# Brain-wide functional connectome analysis of 40,000 individuals reveals brain networks that show aging effects in older adults

**DOI:** 10.1101/2024.05.17.594743

**Authors:** Yezhi Pan, Chuan Bi, Peter Kochunov, Michelle Shardell, J. Carson Smith, Rozalina G. McCoy, Zhenyao Ye, Jiaao Yu, Tong Lu, Yifan Yang, Hwiyoung Lee, Song Liu, Si Gao, Yizhou Ma, Yiran Li, Chixiang Chen, Tianzhou Ma, Ze Wang, Thomas Nichols, L. Elliot Hong, Shuo Chen

## Abstract

The functional connectome changes with aging. We systematically evaluated aging related alterations in the functional connectome using a whole-brain connectome network analysis in 39,675 participants in UK Biobank project. We used adaptive dense network discovery tools to identify networks directly associated with aging from resting-state fMRI data. We replicated our findings in 499 participants from the Lifespan Human Connectome Project in Aging study. The results consistently revealed two motor-related subnetworks (both permutation test p-values <0.001) that showed a decline in resting-state functional connectivity (rsFC) with increasing age. The first network primarily comprises sensorimotor and dorsal/ventral attention regions from precentral gyrus, postcentral gyrus, superior temporal gyrus, and insular gyrus, while the second network is exclusively composed of basal ganglia regions, namely the caudate, putamen, and globus pallidus. Path analysis indicates that white matter fractional anisotropy mediates 19.6% (p<0.001, 95% CI [7.6% 36.0%]) and 11.5% (p<0.001, 95% CI [6.3% 17.0%]) of the age-related decrease in both networks, respectively. The total volume of white matter hyperintensity mediates 32.1% (p<0.001, 95% CI [16.8% 53.0%]) of the aging-related effect on rsFC in the first subnetwork.

## 1 Introduction

Age affects the intrinsic organization of the brain (Damoiseaux, 2017). The alterations in functional and structural connectivity are often linked to upstream and downstream disease processes in major neurodegenerative illnesses (Pievani et al., 2014). Functional connectivity (FC) refers to the temporal correlations in neural activity between brain regions, relying on the underlying anatomical connections established by structural connectivity and providing insights into the integration of brain networks (E. Bullmore & Sporns, 2009; Friston et al., 1993). It is hypothesized that functional brain organization changes precede the structural changes (Jack et al., 2010). Therefore, understanding the pattern of FC change in aging may help identify biomarkers for tracking early disease progression and informing interventions for cognitive health. Among the functional neuroimaging techniques that allow researchers to study human brains in vivo, resting-state functional magnetic resonance imaging (rfMRI) has become a widely used tool due to the discovery of the blood oxygen level-dependent (BOLD) signal, which measures temporal correlations across different brain regions and are robustly correlated during the resting state, hence enabling non-invasive mapping of functional organization of human brains in the absence of specific tasks or stimuli (Attwell et al., 2010; Biswal et al., 1995; Fox & Raichle, 2007; Logothetis, 2002).

Previous rfMRI studies show an overall picture of age-related reduction of resting-state functional connectivity (rsFC) *within* a few higher-order brain networks, including the default mode network (DMN), salience network (SN), cognitive control network (CCN), and dorsal attention network (DAN) (Andrews-Hanna et al., 2007; Damoiseaux et al., 2008; Ferreira et al., 2016; Ferreira & Busatto, 2013; Geerligs et al., 2015; Grady et al., 2016; Ng et al., 2016; Zonneveld et al., 2019). Interestingly, increase of rsFC *between* networks and decreased segregation with advancing age were also commonly reported(Andrews-Hanna et al., 2007; Chan et al., 2014; Ferreira et al., 2016; Geerligs et al., 2015; Grady et al., 2016). Meanwhile, motor and subcortical networks showed more conflicting results (Deery et al., 2023; Ferreira & Busatto, 2013). Some evidence showed that the FC within sensorimotor or somatomotor network declines with aging (Bernard et al., 2013; Jockwitz & Caspers, 2021; Zonneveld et al., 2019); some, on the other hand, indicated an increase within these regions or no significant changes (Cao et al., 2014; Geerligs et al., 2015; King et al., 2018; Song et al., 2014). The age-related alterations in rsFC, both *within* and *between* networks, have been observed in studies related to cognitive performance (e.g., conditions marked by cognitive decline such as Alzheimer’s disease) and motor ability (Andrews-Hanna et al., 2007; Ferreira & Busatto, 2013; Lin et al., 2018; Szewczyk-Krolikowski et al., 2014; Yoshimura et al., 2020). Additionally, some investigations have showcased the promising prospect of predicting cognitive capabilities through comprehensive whole-brain FC (Finn et al., 2015; Rosenberg et al., 2016; Shen et al., 2017). Therefore, elucidating the systematic age-related rsFC changes could offer exciting opportunities for understanding the neural processes underlying the decline in both cognitive and motor performance observed during the natural aging trajectory.

Network analysis of age-related rsFC is commonly used to study the complex connectome variables. Existing studies typically adopted predefined functional networks (Nichols et al., 2017; Yeo et al., 2011) to examine the impact of aging on FC. However, connections that show aging effects may not be fully constrained within a predefined functional network. In practice, age-related edges (i.e., functional connections between brain region pairs) often involve nodes (i.e., brain regions) from two distinct predefined functional networks. The age-related edges may also consolidate into previously unknown and age-related subnetworks. Therefore, relying solely on predefined networks to study the age effect could consequently lead to increased risks of i) low sensitivity and inflated type II error by missing age-related edges not in predefined networks; and ii) increased false positive findings by claiming that a predefined network is age-related when only a small proportion of intra-network edges are age-related.

To address this challenge, we combine data-driven network analysis with predefined networks to investigate age-related connectome patterns. We first employ data-driven network analysis methods to examine whether age influences any rsFC edges and whether the rsFC edges with age-related differences can consolidate into organized *dense* subnetworks (Bullmore & Bassett, 2011; Wu et al., 2022). We define an age-related subnetwork as *dense* when the proportion of age-related intra-network connections (i.e., network density) is high (Tsourakakis et al., 2013). Since age-related subnetworks extracted by data-driven methods are not constrained by predefined networks, they may consist of brain regions from multiple predefined networks. Therefore, we further identify (parts of) predefined networks corresponding to each age-related subnetwork extracted through data-driven network analysis. This approach enables us to simultaneously: i) improve sensitivity by capturing age-related edges from both inter- and intra-predefined networks; ii) reduce the false positive error rate through shrinkage (i.e., *densification*); and iii) yield interpretable network findings by linking with predefined networks. The probability of a data-driven age-related subnetwork being false positive is exponentially determined by its size and *density*. The probability approaches zero for a subnetwork of 10 nodes and a density of 50% (Chen et al., 2023). However, interpreting data-driven age-related subnetworks may not be straightforward as they can consist of nodes from multiple networks. Hence, identifying predefined subnetworks within each data-driven network can assist understanding the systematic effects of aging.

In the present study, we investigated age-related rsFC differences across the entire brain leveraging a large sample, the UK Biobank (UKB) cohort (Sudlow et al., 2015) (n=39,675). We modeled the relationship between measured rsFC and age, and then adopted an adaptive dense subnetwork extraction procedure to to maximize the coverage of age-related rsFC edges throughout the entire brain within subnetworks predominantly composed of these edges (Tsourakakis et al., 2013; Wu et al., 2022) (see Figure 1A). Extracted subnetworks were mapped onto Yeo’s 7 resting-state networks for functional interpretation (Yeo et al., 2011). To ensure reproducibility, we further replicated the analysis with an independent validation sample that includes 499 participants from the Lifespan Human Connectome Project in Aging (HCP-A) cohort (Bookheimer et al., 2019). The utilization of two large population-based samples provides compelling evidence for the robustness of the findings.

**Figure 1.**
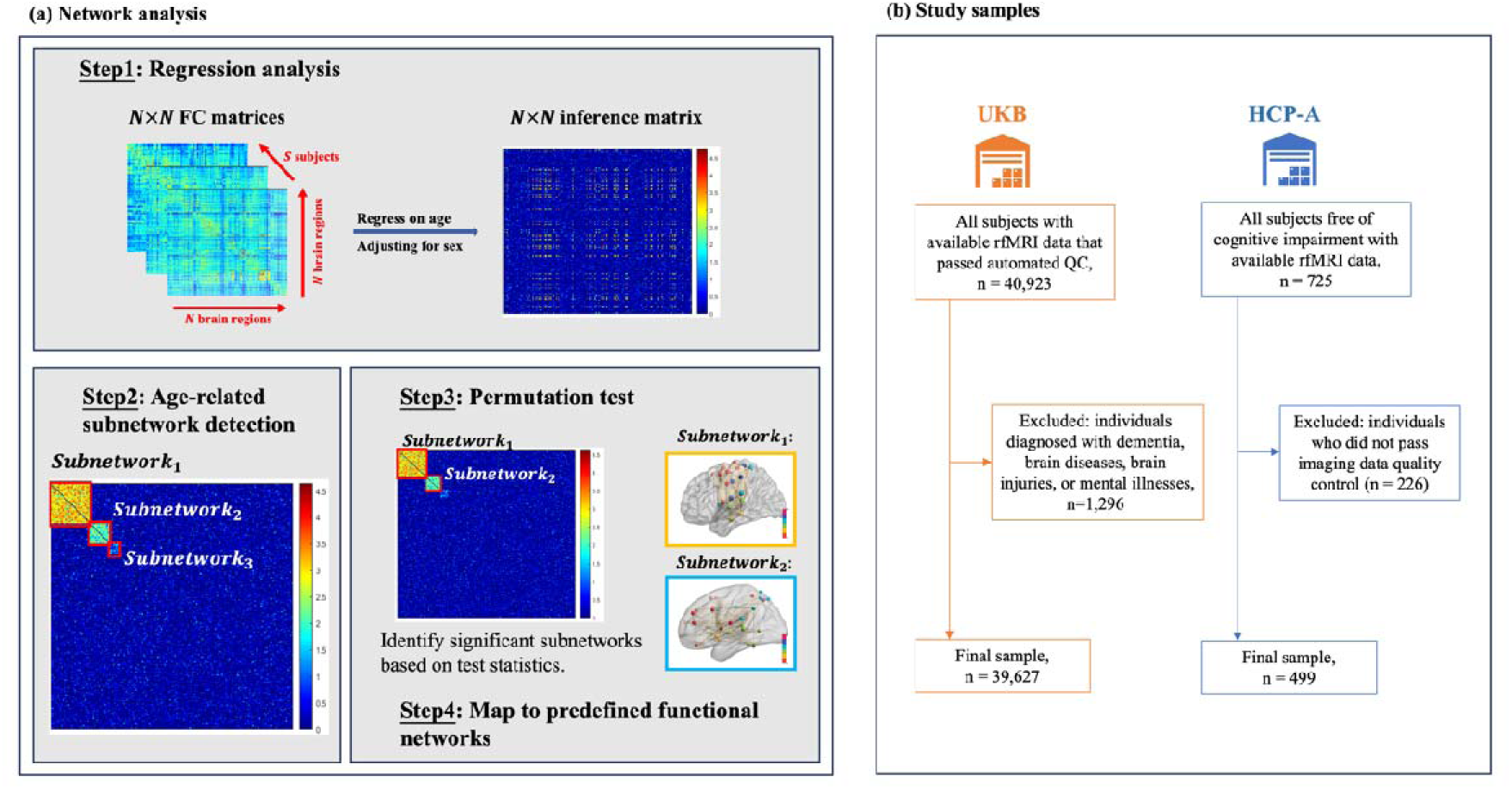
Study design and cohort flow chart. Panel (a) illustrates the data-driven network analysis design in this study. Panel (b) presents the cohort inclusion flowchart.

## 2 Methods

### 2.1 Study samples

We utilized two independent population-based cohorts. The first cohort is the UK Biobank (Sudlow et al., 2015) (UKB, http://www.ukbiobank.ac.uk/), a large prospective study with approximately 500k participants aged 40-69 years at recruitment between 2006-2010 across 22 assessment centers in the UK. Ethical approval for the UKB study was obtained from the National Information Governance Board for Health and Social Care and the National Health Service North West Multicenter Research Ethics Committee (REC reference 21/NW/0157). Written informed consent was obtained from all UKB participants. A total of ∼100k participants underwent brain magnetic resonance imaging (MRI) assessments(Alfaro-Almagro et al., 2018; Miller et al., 2016). In our analysis, we utilized resting-state functional MRI (rfMRI) data from the v1.8 December 2020 release that measured ∼43k participants starting from 2014. The second cohort that served as the validation set is the Lifespan Human Connectome Project in Aging (Bookheimer et al., 2019) (HCP-A, https://www.humanconnectome.org/study/hcp-lifespan-aging). HCP-A is a large sample comprising approximately 1,200 healthy adults aged 36-100+ years, providing extensive data on structural and functional connectivity, with a specific focus on factors influenced by advanced aging. For our study, we utilized the HCP-A rfMRI data from the 2.0 release, consisting of 725 subjects who underwent imaging assessment sessions starting from 2019.

To specifically investigate the effects of aging, participants in the UKB cohort with various forms of dementia, brain diseases, brain injuries, and mental illnesses (n=1,248) were excluded based on ICD-10 codes (refer to Table S1 in the supplementary material for the complete list of excluded conditions). The initial HCP-A cohort consisted of individuals who had not been diagnosed with pathological causes of cognitive decline, and consequently, no exclusion criteria were applied. Figure 1B shows the cohort flowchart.

### 2.2 rfMRI preprocessing

The details of UKB rfMRI data acquisition parameters and artefact removal procedures were described in SI.1 in the supplementary material and elsewhere (Alfaro-Almagro et al., 2018). The downloaded pre-processed rfMRI data were registered to a 2mm MNI152-template (Grabner et al., 2006) using FSL (the FMRIB Software Library) (Jenkinson et al., 2012), in order to normalize the brain images to a standard space and discard non-brain regions. We then used AFNI (Analysis of Functional Neurolmages) (Cox & Hyde, 1997) to extract time series from the normalized rfMRI data for each of 246 ROIs based on The Human Brainnetome Atlas (Fan et al., 2016).

For HCP-A, we aligned the preprocessed rfMRI data across subjects using MSMAll multi-modal surface registration, registered the data to a 2mm MNI152-template using FSL, and extracted time series from 246 Human Brainnetome ROIs using AFNI. Besides the standard quality control during the HCP-A rfMRI data acquisition, we further performed quality control after downloading the preprocessed data and retained a total of 499 participants whose rsFC data had a low missing ratio and an acceptable variation (refer to SI.1.4 in the supplementary material for definition). Missing rsFC data were imputed with the mean rsFC value.

### 2.4 Imaging confounding measures

Previous studies have suggested the adjustment of several potential imaging confounders when conducting research on imaging derived phenotypes (Elliott et al., 2018). Therefore, we controlled for variables that addressed diverse dimensions of imaging confounds during the rsfMRI session, which had the potential to introduce bias into the imaging outcomes. In the UKB, we included the following variables as covariates: 1) mean head motion, quantified through the mean displacement (mm) between consecutive time points across all regions (data-field 25741); 2) scanner lateral brain position, the X-coordinate (left-right) of the center of the brain mask within the scanner (data-field 25756); 3) scanner transverse brain position, the Y-coordinate (front-back) of the back of the brain mask within the scanner (data-field 25757); 4) scanner longitudinal brain position, the Z-coordinate of the center of the brain mask within the scanner (data-field 25758); and 5) scanner table position, the Z-coordinate of the coil within the scanner. The latter four covariates were a set of positioning variables that accounted for variations in the precise head and coil placement within the scanner across different scanned individuals.

### 2.5 Connectivity analysis

Pearson’s correlation of the rfMRI timeseries between each pair of the 246 Human Brainnetome Atlas ROIs was used to quantify the rsFC. This resulted in the quantification of subject-specific rsFC, yielding a total of 30,135 edges for each subject *s*, where *s*= 1, …, *s*; with *s* = 39,675 in UKB and *s* = 499 in HCP-A. The rsFC was denoted as a weighted adjacency matrix Y∈ *R*^*N*×*N*^, where each entry 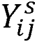 represents the connection strength between the i-th and j-th ROIs. A graph model *G*= {*V*, *E*} was then adopted to describe the topological structure of the brain connectome, with *V* denoting the set of ROIs and *E* denoting the set of rsFC edges within and between ROIs. The data-driven brain connectome analysis involves three main steps: 1) regression analysis on individual rsFC edges, 2) extraction of subnetworks, and 3) statistical inference on the detected subnetworks. These analyses are conducted separately for the UKB and HCP-A cohorts. Lastly, we identified predefined networks within each data-driven subnetwork.

#### 2.5.1 Regression analysis

In the first step, a generalized matrix response regression model was applied to the whole-brain connectome matrix **Y^s^**∈ *R*^*N*×*N*^to determine the age-associated changes in each of the FC edges, adjusting for sex and imaging confounding measures:

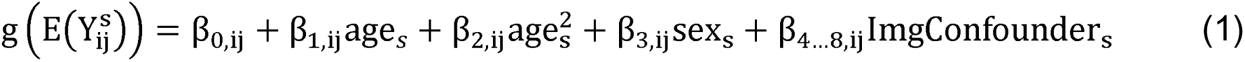

where *g* is the identity link function for continuous connectivity measures and logistic link function for binary connectivity measures. The mass-univariate testing thus yielded an *N*×*N* weighted inference matrix **w** associated with β_1_, where each element *w_ij_* denotes the association of age and FC between the i-th and j-th ROIs. In addition, the quadratic term of age was included only if it was statistically significant with a non-trivial effect size (a partial r^2^ of 1%). We used both -log_10_(*p_ij_*) values and *t_ij_* statistics to characterize the strength and magnitude of the associations. For simplicity, we denoted β*_ij_* as the regression coefficient associated with age in the following analysis.

#### 2.5.2 Subnetwork extraction

We performed age-related subnetwork extraction and inference based on *dense* subnetwork detection methods developed recently(Chen et al., 2023; Wu et al., 2022). In these models, we consider edges e_ij_ ∈ E with β*_ij_* ≠ 0 primarily concentrated within dense subgraphs *G_c_*⊂*G* rather than uniformly distributed throughout the whole-brain connectome G. Let *G_c_*⊂*G* denote a subnetwork comprising |*V_c_*| ROIs and 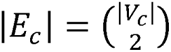 FC edges, such that *G_c_* is associated with age if 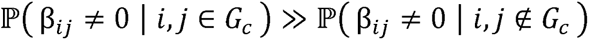. These identified subgraphs, *G_c_*, signify age-related connectome patterns. Herein, we used adaptive dense subgraph extraction algorithms(Tsourakakis et al., 2013) to extract 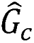, which is an estimate of *G_c_* from w.

#### 2.5.3 Permutation test

For each extracted age-related subgraph 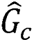, we evaluated its statistical significance through a permutation test with subgraph-tailored test statistics that control the family-wise error rate(Chen et al., 2023). This procedure was carried out to derive a p-value for each extracted subnetwork. Specifically, we randomized the subjects’ ages and performed regression as per Equation 1. This yielded age-related p-values, which together formed the inference matrix **w**^perm^. After applying the subnetwork extraction algorithm, we designated the test statistic *T_k_* for the k-th iteration of the permutation test as the maximum value of the test statistics evaluated for each extracted subnetwork, 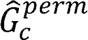. Following M iterations of the permutation test, the p-value associated with 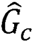 was approximated as the percentile rank of 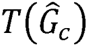 within the set *T*_1_*, T*_2_*,…, T_M_*. For a more detailed explanation of the test statistics formulation, please refer to another work (Chen et al., 2023).

### 2.6 Path analysis

To test if brain white matter microstructure changes or white matter lesion could explain the age-related rsFC declines, we modeled two paths for each subnetwork using linear structural equation models: 1) age ➔ white matter FA ➔ rsFC, 2) age ➔ white matter hyperintensities ➔ rsFC. The FA (UKB data-field 25056) was averaged across the whole-brain for each participant. The total volume of white matter hyperintensities (UKB data-field 25781) was log-transformed. rsFC was averaged across each extracted subnetwork. All models were adjusted for sex.

To evaluate the influence of age-related declines in functional connectivity on cognitive functions, we used linear regression to establish the association between the mean rsFC of both subnetworks and a general cognitive score, while controlling for age. The general cognitive score represented a latent construct of general intelligence, capturing approximately 70% of the total variance across nine cognitive assessments corresponding to seven distinct domains, namely processing speed, perceptual reasoning, visuospatial learning and memory, cognitive flexibility, executive function and planning, working memory, and fluid intelligence. Prior to conducting the analysis, a quality control procedure was executed on the cognitive data (as detailed in Table S3 in the supplementary material). Missing data were imputed using the predictive mean matching (PMM) method. This composite cognitive score serves as a comprehensive metric for assessing overall cognitive performance.

## 3 Results

### 3.1 Participant demographics

The mean age of UKB participants was 63.9 years (range 44-82, standard deviation [SD] 7.7); in contrast, HCP-A had a lower mean, but wider range of ages (mean 60.2 years, range 36-100, SD 15.9). In the UKB cohort, 54.4% of participants were female, while in the HCP-A cohort, the percentage of females was 58.5%.

### 3.2 Age-rsFC association

A heterogeneous pattern of age-related rsFC changes throughout the entire brain was observed (Figure 2). These changes encompassed a mixture of increase and decrease in rsFC. T-statistics of the aging effect on rsFC in both cohorts showed a negative-centering and bell-shaped distribution, suggesting a predominance of age-related decreases rather than increases. The resulting 30,135 t-statistics and -log_10_ transformed p-values associated with the main effect of age were formed into a 246 x 246 inference matrices for each of the two cohorts respectively.

**Figure 2.**
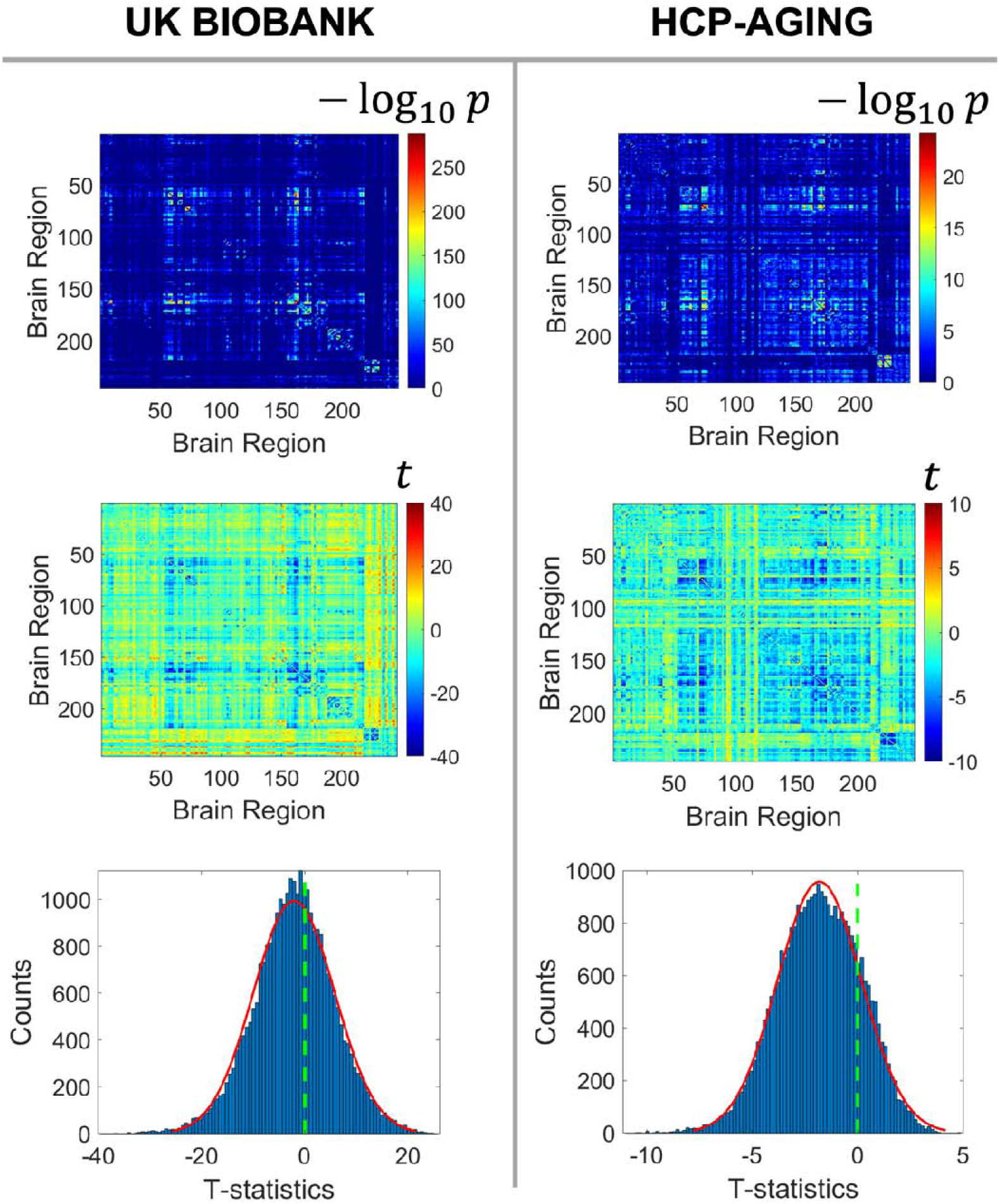
Pattern of age-related rsFC change. The top row shows the inference matrices of -log_10_(p-values) indicating the significance of age-related effects on individual rsFC edges. Higher values (red colors) denote greater significance. The middle row displays the inference matrices of t-statistics that represents the effect size and direction of age-related effects on individual rsFC edges. Negative values (blue colors) indicate age-related declines; positive values (red colors) mean age-related increases. The third row demonstrates the distribution of the t-statistics of the effect of aging on all individual edges. The prevalence of age-related declines in rsFC shift both histograms to the left.

In Figure 2, it appeared that certain latent subnetworks may be systematically influenced by aging, that is, a significant proportion of intra-network edges shows aging effects while only a minor fraction of connections beyond the networks are age-related. Identifying these subnetworks has the potential to enhance our understanding of how aging impacts the brain’s connectome at a network level. Nevertheless, individual age-related connections alone are insufficient to reveal these brain subnetworks. Therefore, it is imperative to employ a data-driven method to extract subnetworks capable of capturing the underlying structure of age-related differences in rsFC.

### 3.3 Subnetworks showing aging effects

The clique-forming regions that show age-related decreases were identified using an adaptive dense subnetwork extraction method (Figure 3). We focused on characterizing rsFC decreases in the following subnetwork analysis. In UKB, the first subnetwork (permutation p-value<0.001) consisted of 59 brain regions. Mapped onto Yeo’s 7 resting-state network, the first dense subnetwork contains 31 regions from sensorimotor network and 13 from ventral and dorsal attention network. The second subnetwork (permutation p-value<0.001) comprised 9 brain regions exclusively from the basal ganglia. Similarly, in HCP-A, the first subnetwork (permutation p-value<0.001) contained 55 brain regions, with 27 regions from the sensorimotor network and 24 regions from ventral and dorsal attention network. The second subnetwork (permutation p-value<0.001) from HCP-A also consisted of 10 brain regions solely from the basal ganglia. Since the sample size of HCP-A is smaller, the noise level of statistical inference (e.g., -log_10_(p) is higher (Figure 3B).

**Figure 3.**
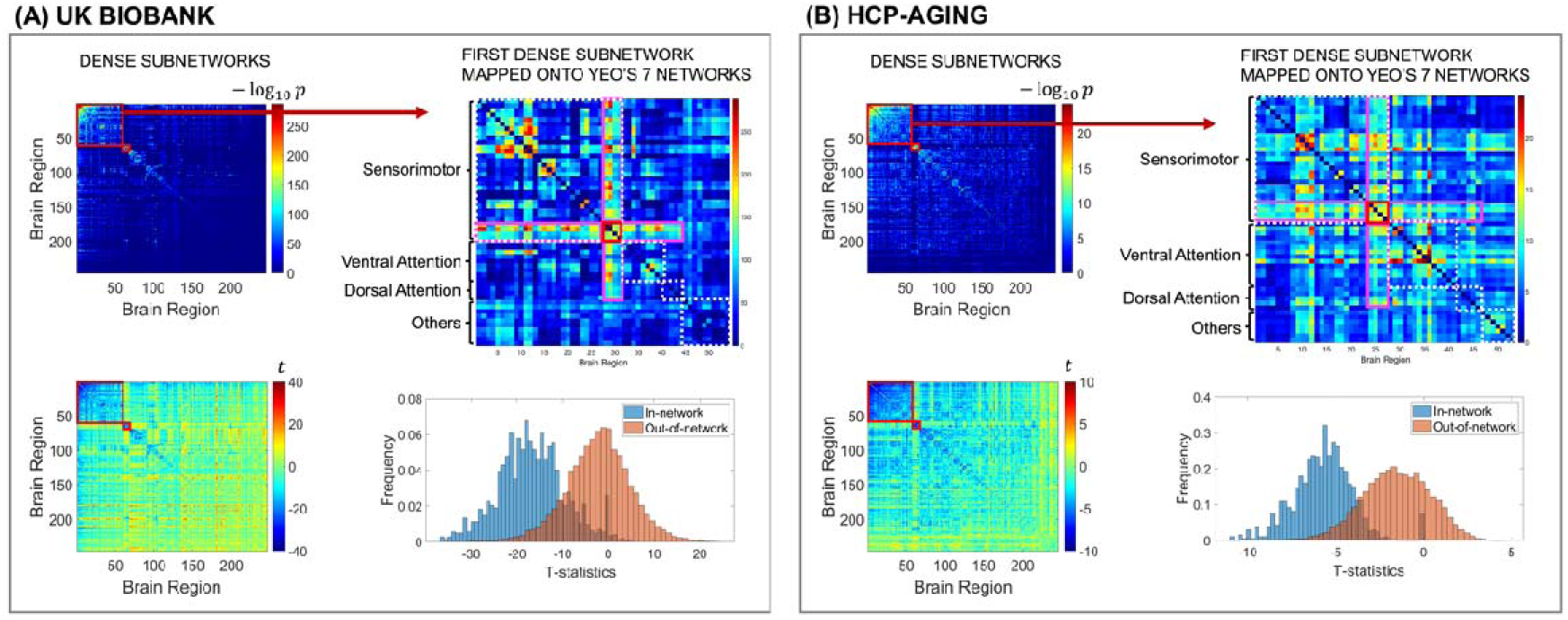
Subnetworks of age-related rsFC differences. Figures on the left in both panels display the dense subnetworks extracted using the data-driven subnetwork detection method, highlighted with red boxes. Each matrix element is -log_10_(p-values) (top left) and corresponding t-statistic (bottom left) obtained from association analysis between each rsFC edge and age. The top right figures in both panels represent the mapping of the large sensorimotor-and-attention-related subnetwork onto Yeo’s 7 resting-state functional networks. The red boxed regions are the hubs consisting of hypergranular insula and dorsal granular insula that have decreased rsFC with the all other sensorimotor-and-attention-related regions in the subnetwork. The bottom right figures in both panels present the systematic differences between the t-statistics within and beyond the extracted subnetworks.

#### 3.3.1 Replicable subnetworks by UKB and HCP-A

The data-driven age-related subnetworks identified in UKB and HCP-A are highly consistent. The sensorimotor-and-attention-related subnetworks from both cohorts had an overlap of 40 brain regions. Within the overlap, 24 distinct regions were ascribed to the sensorimotor network, constituting approximately 73% of all regions characterized as being pertinent to sensorimotor functions. Anatomically, this sensorimotor-and-attention-related subnetwork included the majority (containing ≥ 50% of the total regions in the gyrus) of the precentral gyrus, postcentral gyrus, superior temporal gyrus, paracentral lobule, and insular gyrus - remarkably close to the central sulcus. Furthermore, the secondary subnetworks extracted from both cohorts have an overlap of 9 regions exclusively within the basal ganglia, encompassing 9 out of the total 12 basal ganglia regions, including bilateral putamen, bilateral globus pallidus, bilateral dorsal caudate, and left ventral caudate. The results suggested a systematic age-related decrease in rsFC within the sensorimotor network, dorsal/ventral attention network, and the basal ganglia. The full list of brain regions that showed age-related decreases in rsFC can be found in Table S2 in the supplementary material. Moreover, 4 regions from the insula (bilateral hypergranular insula and bilateral dorsal granular insula) consistently functioned as a hub in the age-related subnetworks. These 4 insula regions showed an age-related decreased connection with all regions from the sensorimotor and dorsal/ventral attention networks (Figure 3). Figure 4 shows the 3-D axial and sagittal representations of the intersection of the subnetworks identified from both UKB and HCP-A.

**Figure 4.**
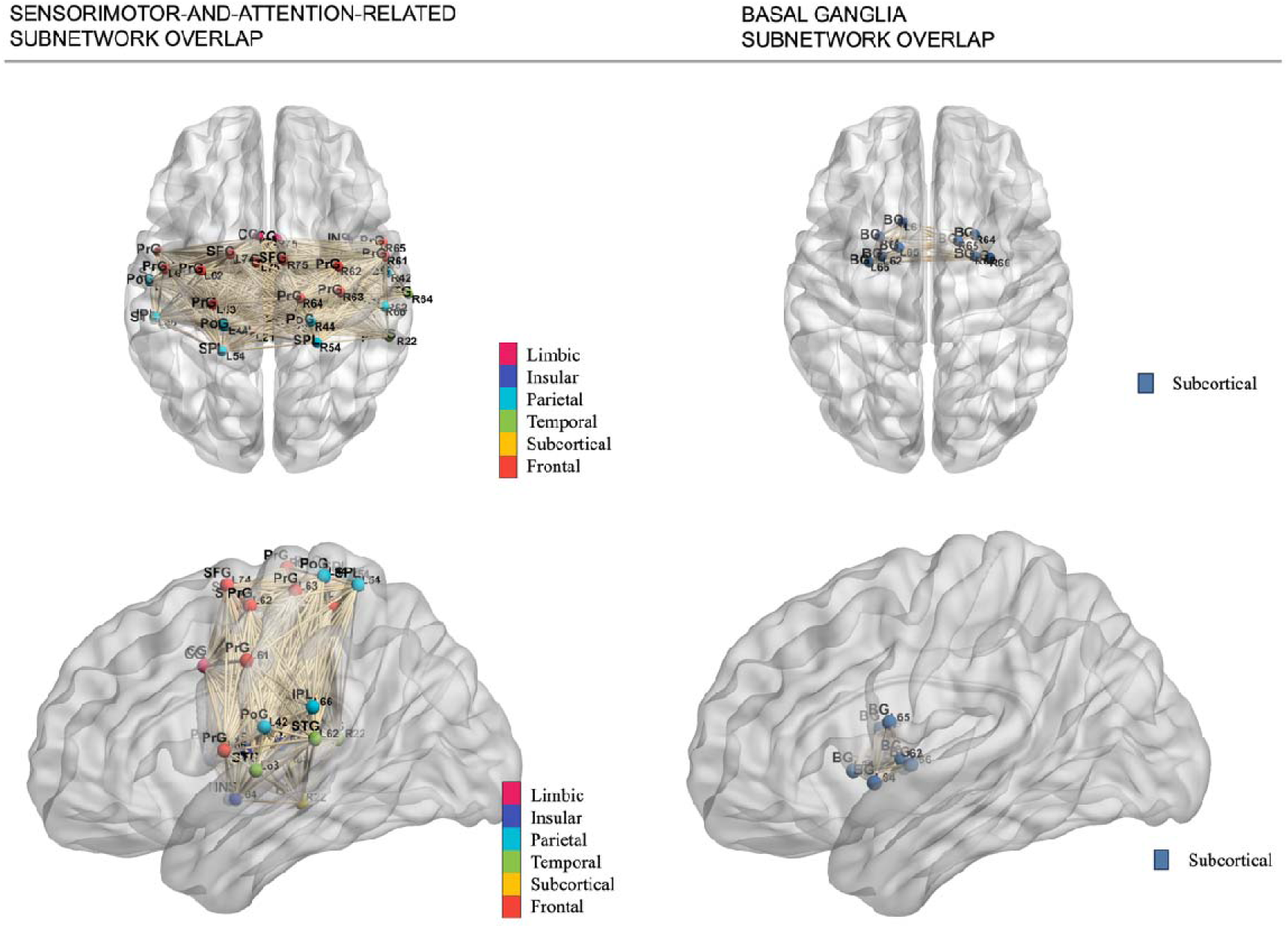
3-D representation of the intersection of extracted age-related connectomic subnetworks. The left row demonstrates the axial and sagittal view of the intersection of the sensorimotor-and-attention-related subnetwork extracted using the data-driven method. The right row shows the axial and sagittal view of the intersection of the basal ganglia subnetwork. Node colors are labeled by cerebral cortex lobes. The 3-D demonstration of individual extracted subnetworks from both UKB and HCP-A are displayed in Figure S3 in the supplementary material.

Additionally, we performed a sensitivity analysis to assess the robustness of the primary analysis results. We conducted the same set of analyses on the entire UKB cohort that passed imaging data quality control (n=40,923), regardless of health status, including participants with neurological diseases, brain injuries, or mental illnesses (n=1,248) who were excluded from the primary analysis. The patterns of age-related declines in rsFC showed negligible variations as compared to the primary result (Figure S2 in the supplementary material). We also conducted a subgroup analysis based on the biological variable of sex. Our analysis revealed no systematic differences in the patterns of rsFC changes between sexes, as illustrated in Figure S1 in the supplementary material.

### 3.5 Structural changes related to age-related functional declines by path analysis

Linear structural equation modeling analysis showed that in the sensorimotor- and-attention-related subnetwork, 19.6% (p<0.001, 95% CI [7.6% 36.0%]) of the total effect of aging on rsFC is mediated through FA, while in the basal ganglia subnetwork, this proportion is 11.5% (p<0.001, 95% CI [6.3% 17.0%]). Moreover, in the sensorimotor-and-attention-related subnetwork, a substantial 32.1% (p<0.001, 95% CI [16.8% 53.0%]) of the aging-related effect on rsFC is mediated by the total volume of white matter hyperintensities, whereas the mediation effect of white matter hyperintensities was found to be statistically insignificant in the basal ganglia subnetwork (model results shown in Figure S4 in the supplementary material). To further understand how the changes in functional connectivity may be linked to differences in cognitive functioning, we modeled the relationship between the average rsFC of each subnetwork and the general cognitive score. The results showed that one unit decrease of rsFC in the sensorimotor-and-attention-related subnetwork is associated with 0.38 unit (p<0.001, 95% CI [0.28 0.49]) decrease of cognitive score, while one unit decrease of rsFC in the basal ganglia subnetwork is associated with 0.13 (p<0.001, 95% CI [0.06 0.19]) unit decrease of cognitive score.

## 4 Discussion

In this population-based study, we used rfMRI data from two large independent cohorts to investigate the brain-wide rsFC showing age-related differences and extracted functional networks exhibiting heightened susceptibility to the aging process. We employed a novel data-driven network analysis method to improve the specificity of predefined network analysis. The neurofunctional findings provided compelling evidence of a significant decrease in rsFC within the sensorimotor network and within basal ganglia as age advances. Our findings were consistent with the consensus in literature that brain networks experience age-related reorganization changes in modularity (Bassett et al., 2010; Geerligs et al., 2015; Meunier, Achard, et al., 2009; Meunier, Lambiotte, et al., 2009; Song et al., 2014). Modularity refers to the organization of brain regions into distinct modules or communities based on their patterns of connectivity. As the organization of brain networks shifts with age, pre-defined networks may no longer accurately describe the dynamic interactions and connectivity patterns that emerge during the aging process. Therefore, our likelihood-based community detection algorithm was able to provide a more holistic characterization of functional reorganization of aging brains based on the latent pattern. This study contributes empirical evidence that can help reconcile conflicting findings in the past studies and shed light on the underlying mechanisms driving the complex relationship between brain function and the aging process.

Previous studies reported conflicting results on age-related rsFC changes within the sensorimotor network(Jockwitz & Caspers, 2021). Some studies showed an increase of connectivity strength in the sensorimotor regions during aging, especially the left supplementary motor area (leftSMA)(Cao et al., 2014; Seidler et al., 2015; Song et al., 2014; Tomasi & Volkow, 2012). In contrast, certain studies have revealed a notable decline in resting-state functional connectivity (rsFC) within the sensorimotor network as individuals age. This finding was observed either across the entirety of the sensorimotor network or specifically within regions such as the cortico-cerebellar or mid-posterior insula regions that are part of the sensorimotor network(Bernard et al., 2013; He et al., 2017; Zonneveld et al., 2019). Additionally, one study reported no changes observed in rsFC within the somatomotor network(Geerligs et al., 2015). One study found a slight decrease of rsFC in the sensorimotor network before 80 years old followed by a slight increase after 80 years old(Farràs-Permanyer et al., 2019). The current controversy regarding how rsFC in sensorimotor changes in the normal aging process could partly be due to the relatively small sample size of all existing studies, most of them conducted with 50 to 200 subjects. The largest study on age-related rsFC alterations to our knowledge was conducted by Zonneveld et al. with a study sample of 2,878 non-demented subjects(Zonneveld et al., 2019). Their findings were consistent with our results, indicating a significantly reduced rsFC in the sensorimotor network at older age. Similarly, age-related functional changes within the basal ganglia also remain a subject of knowledge gaps in current literature. Our age-related FC reduction findings was consistent with a recent large size resting state fMRI study showing age-related brain entropy increase in the motor cortex (Wang, 2020). Increased entropy indicates higher randomness of the fMRI time courses, which would lead to reduced FC. Our result was also consistent with a study that focused on older adults and revealed a negative association between FC and age(Griffanti et al., 2018), whereas it contradicted studies that focused on development or the entire lifespan(Allen et al., 2011; Solé-Padullés et al., 2016). This suggests that the change of FC within the basal ganglia across the lifespan could be U-shaped instead of linear.

We observed the wide-spread age-related rsFC decrease between dorsal granular/hypergranular insula and the rest of the sensorimotor-and-attention-related subnetwork. Conventionally, insular cytoarchitectonic parcellation is divided into the posterior granular section, mid dysgranular section, and anterior agranular section, while the dorsal and posterior part of the insula contains the highest amount of the granule neurons (Morel et al., 2013; Uddin et al., 2017). The posterior granular section (including dorsal granular and hypergranular insula) was found to be functionally connected with the primary and secondary sensorimotor cortices (Deen et al., 2011). The observed age-related decrease in functional connection may indicate decreased sensorimotor network integration and may be linked to declines in motor coordination and cognitive processes commonly observed in the older adult population. The sensorimotor network integration plays a vital role in both regulating motor control and facilitating the learning process, operating across different levels of the central nervous system(Schwartz, 2016). Research demonstrated that changes in sensorimotor function at the cortical level due to aging were linked to shifts in rsFC rather than structural modifications, and proposed that heightened rsFC could be indicative of improved sensorimotor function, exemplified by enhanced performance in arm-reaching tests measuring gap-detection ability(Yoshimura et al., 2020). Additionally, substantial evidence showed that altered sensorimotor integration is associated with the pathophysiology of neurological disorders and movement-related conditions(Dietz & Sinkjaer, 2007; Patel et al., 2014). For example, decreased functional connectivity within the sensorimotor network was found in Parkinson’s disease(Caspers et al., 2021), reinforcing the significance of intact sensorimotor integration for normal motor function and cognitive processes. Interestingly, sensorimotor attenuation, which refers to reduced brain responses to self-movement compared with external stimuli(Weiskrantz et al., 1971), is not only prevalent in the older adult population but also increases with age(Wolpe et al., 2016). The escalating sensorimotor attenuation with age suggests that the neural circuits responsible for self-generated movement perception become less robust, which may in turn lead to a weakening of the synchronized neural activity and connectivity, contributing to the age-related decline in rsFC within the sensorimotor region. This intricate relationship underscores the complex interplay between age-related changes in neural processing, self-perception, and connectivity alterations, collectively shaping the motor and cognitive changes observed in older adults.

The basal ganglia also harbor nuclei responsible primarily for modulating motor control, as well as learning, executive functions, and emotions. Functional connectivity within the basal ganglia has been reported to be associated with many pathological changes and motor functional changes. Just as observed in the sensorimotor network, individuals diagnosed with Parkinson’s disease exhibited reduced rsFC within the basal ganglia when compared to the control group(Szewczyk-Krolikowski et al., 2014; Tan et al., 2015). The age-related reduction in rsFC might represent a vulnerability or predisposition to such disorders. Specifically, our study revealed a decrease in rsFC within bilateral putamen-caudate and bilateral globus pallidus in the aging process, but no change was observed in bilateral nucleus accumbens. The absence of observable changes in the nucleus accumbens raises intriguing questions about the differential effects of aging on distinct components of the basal ganglia network. The nucleus accumbens, a key component of the ventral striatum, is recognized for its involvement in reward processing, motivation, and reinforcement learning(Shirayama & Chaki, 2006). The lack of observable changes could reflect the unique functional role of the nucleus accumbens compared to other basal ganglia nuclei. The nucleus accumbens is particularly linked to motivational processes and the integration of reward-related information, while the other basal ganglia nuclei, such as the putamen and globus pallidus, are primarily involved in motor control and cognitive functions. The nucleus accumbens’ involvement in reward processing and socioemotional functions could render it less sensitive to the same patterns of connectivity alterations observed in other components of the basal ganglia network. Future research incorporating a multimodal approach and task-based connectivity analysis could provide deeper insights into the distinct effects of aging on various components of the basal ganglia.

The path analysis revealed that in the subnetwork with sensorimotor-related regions, both white matter FA and white matter hyperintensities mediate aging’s impact on the functional connectome. The results suggested that the age-related changes in functional connectivity within sensorimotor regions are influenced by a combination of microstructural white matter changes and the burden of white matter hyperintensities, which are often associated with small vessel disease(Kynast et al., 2018). However, in the subnetwork exclusively composed of basal ganglia regions, FA mediates the age-related decline in rsFC, whereas white matter hyperintensities do not. This could imply a specialization in the aging process for different brain regions. The sensorimotor-related regions might be more vulnerable to vascular burden (white matter hyperintensities). In contrast, the basal ganglia regions could have a different response to aging, where microstructural changes in the white matter are more important determinants of functional decline. Additionally, the impact of these two subnetworks on cognitive outcomes appears to be quite different, with the first network having a threefold greater effect on cognitive performance compared to the second network. It’s possible that changes in the sensorimotor network have broader implications for general cognitive function because these regions play a fundamental role in various motor and sensory processes, which, in turn, are essential for cognitive tasks and overall cognitive health. These path analysis findings highlight the region-specific nature of aging effects in the brain.

### Replicable findings between UKB and HCP

The age-related rsFC patterns identified in the UK Biobank (UKB) data were highly replicable in the HCP-A dataset. This high level of replicability stems from recent advancements in imaging acquisition, preprocessing, quality control, and analytical techniques. Under the multiple testing setting, our data-driven network analysis applied *l*_0_ shrinkage to age-related connections in subnetworks, which improved sensitivity and reduced false positive rate in both the UKB and HCP-A datasets resulted in a substantial overlap in findings. The probability for the overlapped findings being false positive (e.g., random noise) in both datasets is < 10^-16^ .

Compared to existing studies, our present study mainly demonstrated two strengths that significantly contributed to its robustness and scientific value. Firstly, we used a large sample and validated our findings with an independent large cohort, which enhanced the study’s statistical power and increases the generalizability of the findings. Secondly, the study utilized data-driven subnetwork extraction algorithm to enhance the findings from pre-defined network analysis. Age-related changes in rsFC were largely heterogeneous in pre-defined networks, challenging the sufficiency of using average results across the pre-defined networks to draw conclusions, possibly reflecting how networks change with age. Therefore, our methodological choice was advantageous as it allowed for novel identification of specific age-related functional subnetworks based on latent structures present within the data itself. This data-driven approach is particularly valuable in complex systems like the brain, where pre- defined networks may not fully capture the intricacies of functional changes.

In summary, using two large independent samples and a data-driven subnetwork detection method, we found that the rsFC of motor-related networks, including the sensorimotor network and the basal ganglia network, could serve as reliable biomarkers for the aging process. These findings underscore the potential of using these networks as early indicators of age-related cognitive and motor decline. This speculation not only provides insight into the intricacies of neural aging but may also pave the way for developing diagnostic tools and interventions aimed at mitigating the impact of age-related motor function changes.

## Data and Code Availability

Data used in this study are publicly available from the UK Biobank (https://www.ukbiobank.ac.uk/) and the Lifespan Human Connectome Project in Aging study (https://www.humanconnectome.org/study/hcp-lifespan-aging). The code will be made publicly available on GitHub.

## Author Contributions

YP and CB performed formal analysis and wrote the manuscript. SC supervised the project and took the lead in editing the manuscript. PK, MS, JCS, ZY, RGM, JY, TL, YY, HL, SL, SG, YM, YL, CC, TN, LEH, TM, and ZW provided critical feedback and helped to shape the research, analysis, and manuscript.

