## supplementary material for "Brain-wide functional connectome analysis of 40,000 individuals reveals brain networks that show aging effects in older adults"

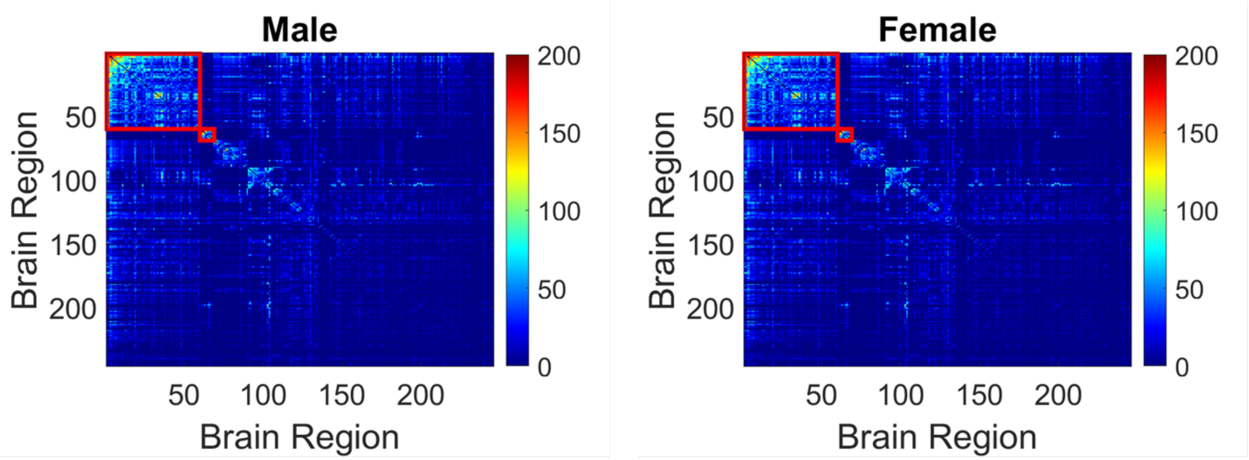


**Figure S1. Age-related subnetworks from sex subgroup analysis.** We performed sex-specific subgroup analysis to assess if there's any difference in age-related functional change patterns between sex. The figures show the subnetworks extracted using the data-driven subnetwork detection method, highlighted within red squares. Each matrix element is the -log10(p-values) obtained from association analysis between each rsFC and age across all subjects within the specific sex group. No difference was observed between the two groups.

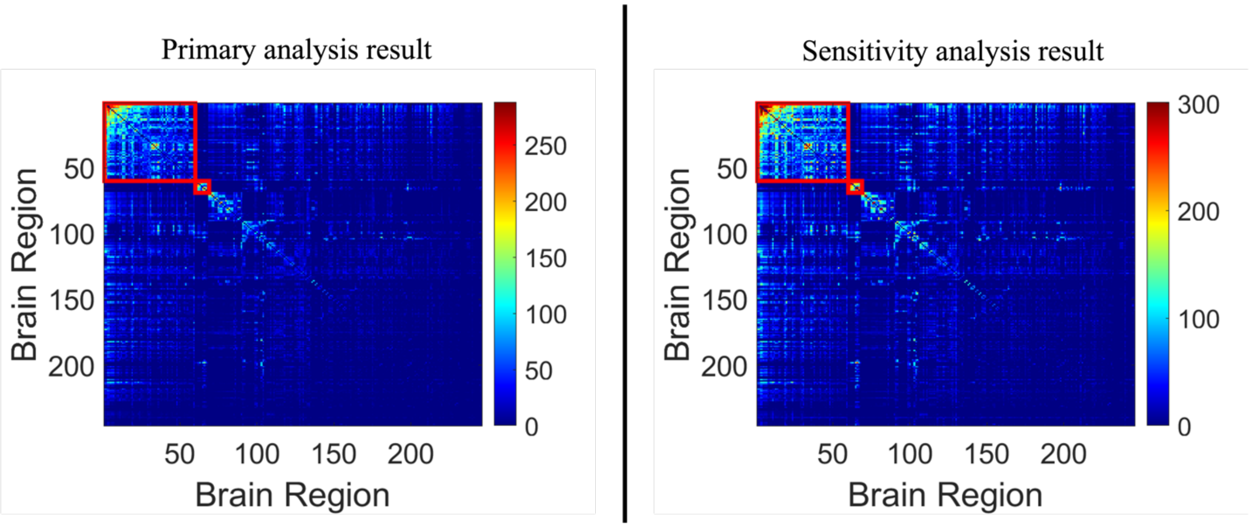


**Figure S2. Sensitivity analysis.** We performed sensitivity analysis on the entire UKB cohort that passed imaging quality check. The sensitivity cohort included 1,248 more subjects than our primary analysis cohort- these were the subjects originally excluded due to their health conditions such as neurological diseases or brain injuries. The highlighted red boxes represent the extracted age-related subnetworks, with each element denoting the -log10(p-values) obtained from association analysis between each rsFC and age across all subjects. The age-related signals become stronger because of a larger sample size, but the difference in the subnetwork patterns is negligible.

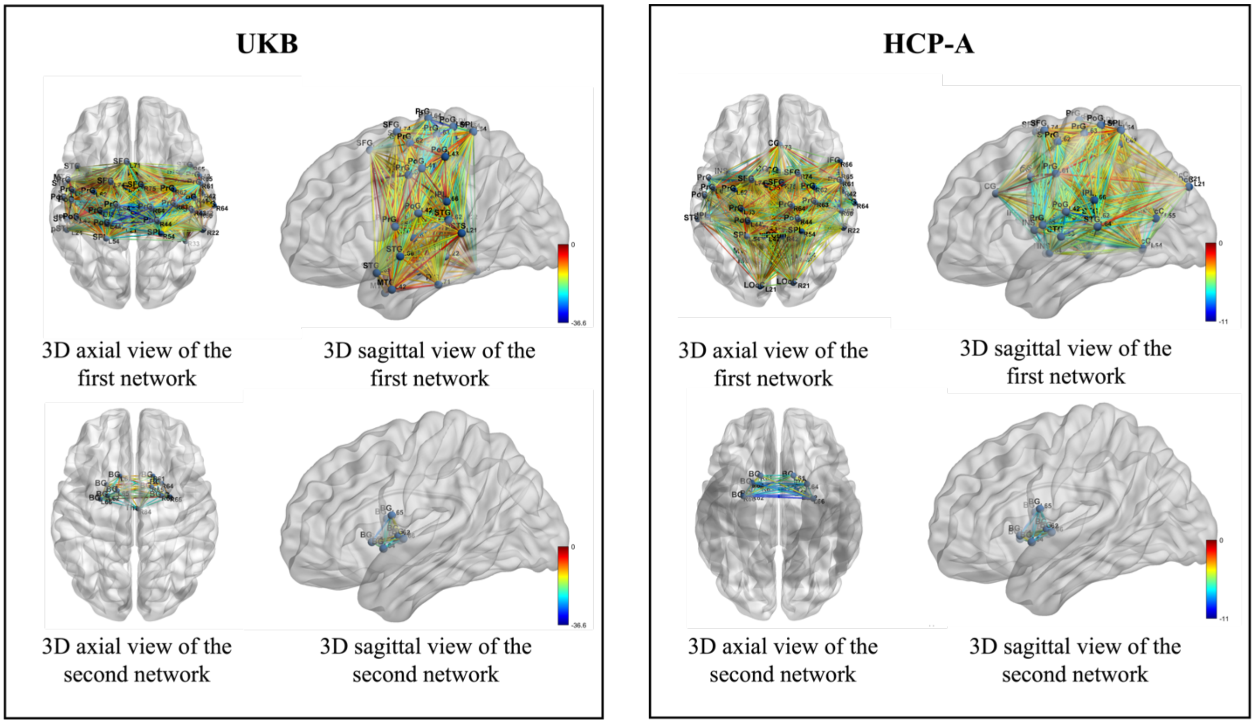


**Figure S3. 3-D demonstration of extracted subnetworks from UKB and HCP-A.** The left panel displays the axial and sagittal views of the first and second age-related subnetworks in UKB, while the right panel presents the corresponding 3-D visualization of age-related subnetworks in HCP-A. Connection colors denote the t-statistics derived from the association analysis between each rsFC and age, with the blue color signifying a larger effect size. Given our focus on age-related rsFC declines, positive associations are not depicted in this representation.

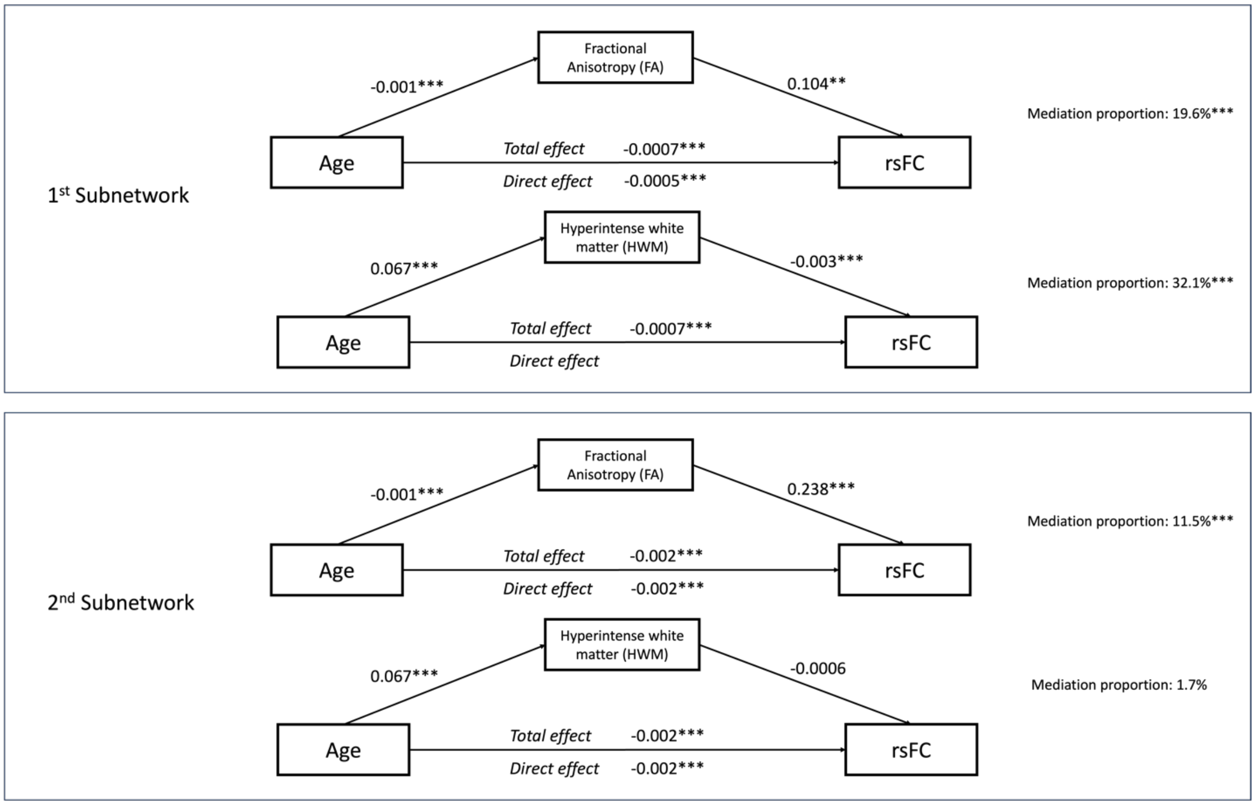


**Figure S4. Mediation effect of structural changes on the age-related functional declines.** We conducted two path analyses for the data-driven age-related subnetworks, respectively. The top model in each panel examines if average FA mediates the effect of aging on the average rsFC of each subnetwork. The bottom model in each panel studies if the total volume of HWM has a mediating effect on age-related functional declines. All models are adjusted for sex. Asterisks indicate significance levels, with one asterisk (*) representing a 0.05 significance level, two asterisks (**) denoting a 0.01 level, and three asterisks (***) indicating a 0.001 level of significance.

### SI.1 rfMRI data preprocessing

#### SI.1.1 rfMRI data acquisition

In UKB, the MRI scans were performed on Siemens Skyra 3-Tesla with a Siemens 32-channel RF receive head coil (protocols available at https://biobank.ctsu.ox.ac.uk/crystal/crystal/docs/brain_mri.pdf). rfMRI data were obtained during a 6min session (490 timepoints, repetition time = 735 ms, echo time = 39 ms, multi-band acceleration 8) with a 2.4mm isotropic resolution and 64 slices.

In HCP-A, the MRI scans were performed on Siemens Prisma 3-Tesla with a Siemens 32-channel Prisma head coil (protocols available at https://www.humanconnectome.org/study/hcp-lifespan-aging/project-protocol/imaging-protocols-hcp-aging). rfMRI data were obtained during a 6min41s-long session (488 timepoints, repetition time = 800 ms, echo time = 37 ms, multi-band acceleration 8) with a 2.0mm isotropic resolution and 72 slices.

#### SI.1.2 rfMRI data preprocessing

Downloaded rfMRI data from UKB underwent echo-planar imaging (EPI) unwarping, gradient distortion correction (GDC), and motion correction using MC-FLIRT (Jenkinson et al., 2002) to reduce interpolation artefacts, and was then FIX-cleaned (Salimi-Khorshidi et al., 2014) to remove structural artefacts.

Downloaded rfMRI data from HCP-A underwent volumetric preprocessing (including GDC, motion correction using FLIRT, slice timing correction, spatial normalization, and spatial smoothing) and surface-based preprocessing (including surface reconstruction, registration to the structural T1w image, and smoothing) (Glasser et al., 2013; Smith et al., 2013). Additionally, multi-run FIX (MR-FIX) was applied to yield better separation of signal and noise components.

#### SI.1.3 UKB rfMRI data quality control

All UKB imaging data was subject to automated quality control based on a standard preprocessing pipeline (Alfaro-Almagro et al., 2018), in which subjects were excluded if their T1-weighted structural scans failed to map to standard space due to missingness, bad head movement, bad field/contrast problem, atypical structure, etc. The full list of quality control measures can be found elsewhere (Alfaro-Almagro et al., 2018).

#### SI.1.4 HCP-A rfMRI data quality control

All HCP-A subjects were subject to quality control measures at various stages, starting with real-time oversight during data acquisition and followed by manual and automated image reviews after acquisition.

We conducted additional quality control measures on the preprocessed HCP-A rfMRI data in response to observed instances of missing data and signal drops in the rfMRI time series. Our approach involved two main steps:

1) We initially excluded all participants whose rsFC data had a missing rate exceeding 5%.

2) To address unexpected signal drops that resulted in extremely high temporal correlations and minimal rsFC variations, we derived a measure to identify participants meeting either of the following criteria:

    a. The ratio of rsFC values larger than 0.9 over all rsFC edges > 0.1 (refer to Figure S5a).
    b. The standard deviation of rsFC values < 0.1 (refer to Figure S5b).

As a result of these quality control procedures, a total of 226 participants (30%) were excluded from the HCP-A cohort.

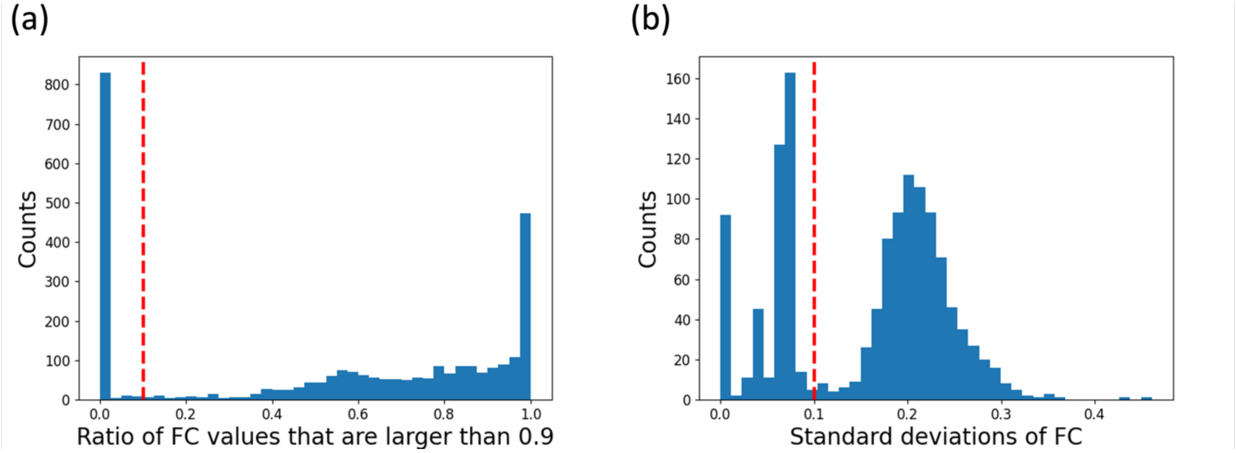


**Figure S5. Measures of variation in the rsFC extracted from HCP-A rfMRI data.** Panel (a) shows a histogram illustrating the ratio of rsFC values exceeding 0.9 across all rsFC edges. The red dashed line at 0.1 represents the threshold we implemented to prevent rsFC with extremely large values from dominating the dataset, which typically occurs because the time series suffer from signal drops due to over-smoothing. Such signal drops in turn can cause the time series to have a correlation (i.e., rsFC) close to 1. This threshold was applied to ensure rsFC sufficient variation. Panel (b) shows the standard deviation of the rsFC data. The red dashed line at 0.1 signifies the cutoff we used to further ensure variation in the rsFC data.

**Table S1. List of ICD-10 codes for various dememtia, neurological disease, and mental illness conditions excluded from the UKB sample**

| ICD-10 Codes | Descriptions |
|---|---|
| A80-A89 | Viral infections of the central nervous system |
| C71 | Malignant neoplasm of brain |
| F00-F09 | Organic, including symptomatic, mental disorders |
| F20-F29 | Schizophrenia, schizotypal and delusional disorders |
| F31 | Bipolar affective disorder |
| F80-F89 | Disorders of psychological development |
| G30-G32 | Other degenerative diseases of the nervous system |
| G35-G37 | Demyelinating diseases of the central nervous system |
| I60-I69 | Cerebrovascular diseases |
| Q00-Q07 | Congenital malformations of the nervous system |
| S00-S09 | Injuries to the head |
| E85 | Amyloidosis |

### Table S2a. Brain regions in the extracted subnetworks from UKB

| Lobe | Gyrus | Left and Right Hemisphere | Label ID.L | Label ID.R | Anatomical and modified Cyto-architectonic descriptions | lh.MNI(X,Y,Z) | rh.MNI(X,Y,Z) |
|---|---|---|---|---|---|---|---|
| Frontal Lobe | SFG, Superior Frontal Gyrus | SFG_L(R)_7_1 | 1 | 2 | *A8m, medial area 8* | -5 ,15, 54 | 7, 16, 54 |
| | | SFG_L(R)_7_2 | 3 | 4 | *A8dl, dorsolateral area 8* | -18, 24, 53 | 22, 26, 51 |
| | | SFG_L(R)_7_3 | 5 | 6 | *A9l, lateral area 9* | -11, 49, 40 | 13, 48, 40 |
| | | SFG_L(R)_7_4 | 7 | 8 | *A6dl, dorsolateral area 6* | -18, -1, 65 | 20, 4, 64 |
| | | SFG_L(R)_7_5 | 9 | 10 | *A6m, medial area 6* | -6, -5, 58 | 7, -4, 60 |
| | | SFG_L(R)_7_6 | 11 | 12 | *A9m,medial area 9* | -5, 36, 38 | 6, 38, 35 |
| | | SFG_L(R)_7_7 | 13 | 14 | *A10m, medial area 10* | -8, 56, 15 | 8, 58, 13 |
| | MFG, Middle Frontal Gyrus | MFG_L(R)_7_1 | 15 | 16 | *A9/46d, dorsal area 9/46* | -27, 43, 31 | 30, 37, 36 |
| | | MFG_L(R)_7_2 | 17 | 18 | *IFJ, inferior frontal junction* | -42, 13, 36 | 42, 11, 39 |
| | | MFG_L(R)_7_3 | 19 | 20 | *A46, area 46* | -28, 56, 12 | 28, 55, 17 |
| | | MFG_L(R)_7_4 | 21 | 22 | *A9/46v, ventral area 9/46* | -41, 41, 16 | 42, 44, 14 |
| | | MFG_L(R)_7_5 | 23 | 24 | *A8vl, ventrolateral area 8* | -33, 23, 45 | 42, 27, 39 |
| | | MFG_L(R)_7_6 | 25 | 26 | *A6vl, ventrolateral area 6* | -32, 4, 55 | 34, 8, 54 |
| | | MFG_L(R)_7_7 | 27 | 28 | *A10l, lateral area10* | -26, 60, -6 | 25, 61, -4 |
| | IFG, Inferior Frontal Gyrus | IFG_L(R)_6_1 | 29 | 30 | *A44d,dorsal area 44* | -46, 13, 24 | 45, 16, 25 |
| | | IFG_L(R)_6_2 | 31 | 32 | *IFS, inferior frontal sulcus* | -47, 32, 14 | 48, 35, 13 |
| | | IFG_L(R)_6_3 | 33 | 34 | *A45c, caudal area 45* | -53, 23, 11 | 54, 24, 12 |
| | | IFG_L(R)_6_4 | 35 | 36 | *A45r, rostral area 45* | -49, 36, -3 | 51, 36, -1 |
| | | IFG_L(R)_6_5 | 37 | 38 | *A44op, opercular area 44* | -39, 23, 4 | 42, 22, 3 |
| | | IFG_L(R)_6_6 | 39 | 40 | *A44v, ventral area 44* | -52, 13, 6 | 54, 14, 11 |
| | OrG, Orbital Gyrus | OrG_L(R)_6_1 | 41 | 42 | *A14m, medial area 14* | -7, 54, -7 | 6, 47, -7 |
| | | OrG_L(R)_6_2 | 43 | 44 | *A12/47o, orbital area 12/47* | -36, 33, -16 | 40, 39, -14 |
| | | OrG_L(R)_6_3 | 45 | 46 | *A11l, lateral area 11* | -23, 38, -18 | 23, 36, -18 |
| | | OrG_L(R)_6_4 | 47 | 48 | *A11m, medial area 11* | -6, 52, -19 | 6, 57, -16 |
| | | OrG_L(R)_6_5 | 49 | 50 | *A13, area 13* | -10, 18, -19 | 9, 20, -19 |
| | | OrG_L(R)_6_6 | 51 | 52 | *A12/47l, lateral area 12/47* | -41, 32, -9 | 42, 31, -9 |
| | PrG, Precentral Gyrus | PrG_L(R)_6_1 | 53 | 54 | *A4hf, area 4(head and face region)* | -49, -8, 39 | 55, -2, 33 |
| | | PrG_L(R)_6_2 | 55 | 56 | *A6cdl, caudal dorsolateral area 6* | -32, -9, 58 | 33, -7, 57 |
| | | PrG_L(R)_6_3 | 57 | 58 | *A4ul, area 4(upper limb region)* | -26, -25, 63 | 34, -19, 59 |
| | | PrG_L(R)_6_4 | 59 | 60 | *A4t, area 4(trunk region)* | -13, -20, 73 | 15, -22, 71 |
| | | PrG_L(R)_6_5 | 61 | 62 | *A4tl, area 4(tongue and larynx region)* | -52, 0, 8 | 54, 4, 9 |
| | | PrG_L(R)_6_6 | 63 | 64 | *A6cvl, caudal ventrolateral area 6* | -49, 5, 30 | 51, 7, 30 |
| | PCL, Paracentral Lobule | PCL_L(R)_2_1 | 65 | 66 | *A1/2/3ll, area1/2/3 (lower limb region)* | -8, -38, 58 | 10, -34, 54 |
| | | PCL_L(R)_2_2 | 67 | 68 | *A4ll, area 4, (lower limb region)* | -4, -23, 61 | 5, -21, 61 |
| Temporal Lobe | STG, Superior Temporal Gyrus | STG_L(R)_6_1 | 69 | 70 | *A38m, medial area 38* | -32, 14, -34 | 31, 15, -34 |
| | | STG_L(R)_6_2 | 71 | 72 | *A41/42, area 41/42* | -54, -32, 12 | 54, -24, 11 |
| | | STG_L(R)_6_3 | 73 | 74 | *TE1.0 and TE1.2* | -50, -11, 1 | 51, -4, -1 |
| | | STG_L(R)_6_4 | 75 | 76 | *A22c, caudal area 22* | -62, -33, 7 | 66, -20, 6 |
| | | STG_L(R)_6_5 | 77 | 78 | *A38l, lateral area 38* | -45, 11, -20 | 47, 12, -20 |
| | | STG_L(R)_6_6 | 79 | 80 | *A22r, rostral area 22* | -55, -3, -10 | 56, -12, -5 |
| | MTG, Middle Temporal Gyrus | MTG_L(R)_4_1 | 81 | 82 | *A21c, caudal area 21* | -65, -30, -12 | 65, -29, -13 |
| | | MTG_L(R)_4_2 | 83 | 84 | *A21r, rostral area 21* | -53, 2, -30 | 51, 6, -32 |
| | | MTG_L(R)_4_3 | 85 | 86 | *A37dl, dorsolateral area37* | -59, -58, 4 | 60, -53, 3 |
| | | MTG_L(R)_4_4 | 87 | 88 | *aSTS, anterior superior temporal sulcus* | -58, -20, -9 | 58, -16, -10 |
| | ITG, Inferior Temporal Gyrus | ITG_L(R)_7_1 | 89 | 90 | *A20iv, intermediate ventral area 20* | -45, -26, -27 | 46, -14, -33 |
| | | ITG_L(R)_7_2 | 91 | 92 | *A37elv, extreme lateroventral area37* | -51, -57, -15 | 53, -52, -18 |
| | | ITG_L(R)_7_3 | 93 | 94 | *A20r, rostral area 20* | -43, -2, -41 | 40, 0, -43 |
| | | ITG_L(R)_7_4 | 95 | 96 | *A20il, intermediate lateral area 20* | -56, -16, -28 | 55, -11, -32 |
| | | ITG_L(R)_7_5 | 97 | 98 | *A37vl, ventrolateral area 37* | -55, -60, -6 | 54, -57, -8 |
| | | ITG_L(R)_7_6 | 99 | 100 | *A20cl, caudolateral of area 20* | -59, -42, -16 | 61, -40, -17 |
| | | ITG_L(R)_7_7 | 101 | 102 | *A20cv, caudoventral of area 20* | -55, -31, -27 | 54, -31, -26 |
| | FuG, Fusiform Gyrus | FuG_L(R)_3_1 | 103 | 104 | *A20rv, rostroventral area 20* | -33, -16, -32 | 33, -15, -34 |
| | | FuG_L(R)_3_2 | 105 | 106 | *A37mv, medioventral area37* | -31, -64, -14 | 31, -62, -14 |
| | | FuG_L(R)_3_3 | 107 | 108 | *A37lv, lateroventral area37* | -42, -51, -17 | 43, -49, -19 |
| | PhG, Parahippocampal Gyrus | PhG_L(R)_6_1 | 109 | 110 | *A35/36r, rostral area 35/36* | -27, -7, -34 | 28, -8, -33 |
| | | PhG_L(R)_6_2 | 111 | 112 | *A35/36c, caudal area 35/36* | -25, -25, -26 | 26, -23, -27 |
| | | PhG_L(R)_6_3 | 113 | 114 | *TL, area TL (lateral PPHC, posterior parahippocampal gyrus)* | -28, -32, -18 | 30, -30, -18 |
| | | PhG_L(R)_6_4 | 115 | 116 | *A28/34, area 28/34 (EC, entorhinal cortex)* | -19, -12, -30 | 19, -10, -30 |
| | | PhG_L(R)_6_5 | 117 | 118 | *TI, area TI(temporal agranular insular cortex)* | -23, 2, -32 | 22, 1, -36 |
| | | PhG_L(R)_6_6 | 119 | 120 | *TH, area TH (medial PPHC)* | -17, -39, -10 | 19, -36, -11 |
| | pSTS, posterior Superior Temporal Sulcus | pSTS_L(R)_2_1 | 121 | 122 | *rpSTS, rostroposterior superior temporal sulcus* | -54, -40, 4 | 53, -37, 3 |
| | | pSTS_L(R)_2_2 | 123 | 124 | *cpSTS, caudoposterior superior temporal sulcus* | -52, -50, 11 | 57, -40, 12 |
| Parietal Lobe | SPL, Superior Parietal Lobule | SPL_L(R)_5_1 | 125 | 126 | *A7r, rostral area 7* | -16, -60, 63 | 19, -57, 65 |
| | | SPL_L(R)_5_2 | 127 | 128 | *A7c, caudal area 7* | -15, -71, 52 | 19, -69, 54 |
| | | SPL_L(R)_5_3 | 129 | 130 | *A5l, lateral area 5* | -33, -47, 50 | 35, -42, 54 |
| | | SPL_L(R)_5_4 | 131 | 132 | *A7pc, postcentral area 7* | -22, -47, 65 | 23, -43, 67 |
| | | SPL_L(R)_5_5 | 133 | 134 | *A7ip, intraparietal area 7(hIP3)* | -27, -59, 54 | 31, -54, 53 |
| | IPL, Inferior Parietal Lobule | IPL_L(R)_6_1 | 135 | 136 | *A39c, caudal area 39(PGp)* | -34, -80, 29 | 45, -71, 20 |
| | | IPL_L(R)_6_2 | 137 | 138 | *A39rd, rostrodorsal area 39(Hip3)* | -38, -61, 46 | 39, -65, 44 |
| | | IPL_L(R)_6_3 | 139 | 140 | *A40rd, rostrodorsal area 40(PFt)* | -51, -33, 42 | 47, -35, 45 |
| | | IPL_L(R)_6_4 | 141 | 142 | *A40c, caudal area 40(PFm)* | -56, -49, 38 | 57, -44, 38 |
| | | IPL_L(R)_6_5 | 143 | 144 | *A39rv, rostroventral area 39(PGa)* | -47, -65, 26 | 53, -54, 25 |
| | | IPL_L(R)_6_6 | 145 | 146 | *A40rv, rostroventral area 40(PFop)* | -53, -31, 23 | 55, -26, 26 |
| | Pcun, Precuneus | PCun_L(R)_4_1 | 147 | 148 | *A7m, medial area 7(PEp)* | -5, -63, 51 | 6, -65, 51 |
| | | PCun_L(R)_4_2 | 149 | 150 | *A5m, medial area 5(PEm)* | -8, -47, 57 | 7, -47, 58 |
| | | PCun_L(R)_4_3 | 151 | 152 | *dmPOS, dorsomedial parietooccipital sulcus(PEr)* | -12, -67, 25 | 16, -64, 25 |
| | | PCun_L(R)_4_4 | 153 | 154 | *A31, area 31 (Lc1)* | -6, -55, 34 | 6, -54, 35 |
| | PoG, Postcentral Gyrus | PoG_L(R)_4_1 | 155 | 156 | *A1/2/3ulhf, area 1/2/3(upper limb, head and face region)* | -50, -16, 43 | 50, -14, 44 |
| | | PoG_L(R)_4_2 | 157 | 158 | *A1/2/3tonIa, area 1/2/3(tongue and larynx region)* | -56, -14, 16 | 56, -10, 15 |
| | | PoG_L(R)_4_3 | 159 | 160 | *A2, area 2* | -46, -30, 50 | 48, -24, 48 |
| | | PoG_L(R)_4_4 | 161 | 162 | *A1/2/3tru, area1/2/3(trunk region)* | -21, -35, 68 | 20, -33, 69 |
| Insular Lobe | INS, Insular Gyrus | INS_L(R)_6_1 | 163 | 164 | *G, hypergranular insula* | -36, -20, 10 | 37, -18, 8 |
| | | INS_L(R)_6_2 | 165 | 166 | *vIa, ventral agranular insula* | -32, 14, -13 | 33, 14, -13 |
| | | INS_L(R)_6_3 | 167 | 168 | *dIa, dorsal agranular insula* | -34, 18, 1 | 36, 18, 1 |
| | | INS_L(R)_6_4 | 169 | 170 | *vId/vIg, ventral dysgranular and granular insula* | -38, -4, -9 | 39, -2, -9 |
| | | INS_L(R)_6_5 | 171 | 172 | *dIg, dorsal granular insula* | -38, -8, 8 | 39, -7, 8 |
| | | INS_L(R)_6_6 | 173 | 174 | *dId, dorsal dysgranular insula* | -38, 5, 5 | 38, 5, 5 |

| | | | | | | | |
|---|---|---|---|---|---|---|---|
| **Limbic Lobe** | **CG, Cingulate Gyrus** | CG_L(R)_7_1 | 175 | 176 | *A23d, dorsal area 23* | -4, -39, 31 | 4, -37, 32 |
| | | CG_L(R)_7_2 | 177 | 178 | *A24rv, rostroventral area 24* | -3, 8, 25 | 5, 22, 12 |
| | | CG_L(R)_7_3 | 179 | 180 | *A32p, pregenual area 32* | -6, 34, 21 | 5, 28, 27 |
| | | CG_L(R)_7_4 | 181 | 182 | *A23v, ventral area 23* | -8, -47, 10 | 9, -44, 11 |
| | | CG_L(R)_7_5 | 183 | 184 | *A24cd, caudodorsal area 24* | -5, 7, 37 | 4, 6, 38 |
| | | CG_L(R)_7_6 | 185 | 186 | *A23c, caudal area 23* | -7, -23, 41 | 6, -20, 40 |
| | | CG_L(R)_7_7 | 187 | 188 | *A32sg, subgenual area 32* | -4, 39, -2 | 5, 41, 6 |
| **Occipital Lobe** | **MVOcC, MedioVentral Occipital Cortex** | MVOcC _L(R)_5_1 | 189 | 190 | *cLinG, caudal lingual gyrus* | -11, -82, -11 | 10, -85, -9 |
| | | MVOcC _L(R)_5_2 | 191 | 192 | *rCunG, rostral cuneus gyrus* | -5, -81, 10 | 7, -76, 11 |
| | | MVOcC _L(R)_5_3 | 193 | 194 | *cCunG, caudal cuneus gyrus* | -6, -94, 1 | 8, -90, 12 |
| | | MVOcC _L(R)_5_4 | 195 | 196 | *rLinG, rostral lingual gyrus* | -17, -60, -6 | 18, -60, -7 |
| | | MVOcC _L(R)_5_5 | 197 | 198 | *vmPOS,ventromedial parietooccipital sulcus* | -13, -68, 12 | 15, -63, 12 |
| | **LOcC, lateral Occipital Cortex** | LOcC_L(R)_4_1 | 199 | 200 | *mOccG, middle occipital gyrus* | -31, -89, 11 | 34, -86, 11 |
| | | LOcC _L(R)_4_2 | 201 | 202 | *V5/MT+, area V5/MT+* | -46, -74, 3 | 48, -70, -1 |
| | | LOcC _L(R)_4_3 | 203 | 204 | *OPC, occipital polar cortex* | -18, -99, 2 | 22, -97, 4 |
| | | LOcC_L(R)_4_4 | 205 | 206 | *iOccG, inferior occipital gyrus* | -30, -88, -12 | 32, -85, -12 |
| | | LOcC _L(R)_2_1 | 207 | 208 | *msOccG, medial superior occipital gyrus* | -11, -88, 31 | 16, -85, 34 |
| | | LOcC _L(R)_2_2 | 209 | 210 | *lsOccG, lateral superior occipital gyrus* | -22, -77, 36 | 29, -75, 36 |
| **Subcortical Nuclei** | **Amyg, Amygdala** | Amyg_L(R)_2_1 | 211 | 212 | *mAmyg, medial amygdala* | -19, -2, -20 | 19, -2, -19 |
| | | Amyg_L(R)_2_2 | 213 | 214 | *lAmyg, lateral amygdala* | -27, -4, -20 | 28, -3, -20 |
| | **Hipp, Hippocampus** | Hipp_L(R)_2_1 | 215 | 216 | *rHipp, rostral hippocampus* | -22, -14, -19 | 22, -12, -20 |
| | | Hipp_L(R)_2_2 | 217 | 218 | *cHipp, caudal hippocampus* | -28, -30, -10 | 29, -27, -10 |
| | **BG, Basal Ganglia** | BG_L(R)_6_1 | 219 | 220 | *vCa, ventral caudate* | -12, 14, 0 | 15, 14, -2 |
| | | BG_L(R)_6_2 | 221 | 222 | *GP, globus pallidus* | -22, -2, 4 | 22, -2, 3 |
| | | BG_L(R)_6_3 | 223 | 224 | *NAC, nucleus accumbens* | -17, 3, -9 | 15, 8, -9 |
| | | BG_L(R)_6_4 | 225 | 226 | *vmPu, ventromedial putamen* | -23, 7, -4 | 22, 8, -1 |
| | | BG_L(R)_6_5 | 227 | 228 | *dCa, dorsal caudate* | -14, 2, 16 | 14, 5, 14 |
| | | BG_L(R)_6_6 | 229 | 230 | *dlPu, dorsolateral putamen* | -28, -5, 2 | 29, -3, 1 |
| | **Tha, Thalamus** | Tha_L(R)_8_1 | 231 | 232 | *mPFtha, medial pre-frontal thalamus* | -7, -12, 5 | 7, -11, 6 |
| | | Tha_L(R)_8_2 | 233 | 234 | *mPMtha, pre-motor thalamus* | -18, -13, 3 | 12, -14, 1 |
| | | Tha_L(R)_8_3 | 235 | 236 | *Stha, sensory thalamus* | -18, -23, 4 | 18, -22, 3 |
| | | Tha_L(R)_8_4 | 237 | 238 | *rTtha, rostral temporal thalamus* | -7, -14, 7 | 3, -13, 5 |
| | | Tha_L(R)_8_5 | 239 | 240 | *PPtha, posterior parietal thalamus* | -16, -24, 6 | 15, -25, 6 |
| | | Tha_L(R)_8_6 | 241 | 242 | *Otha, occipital thalamus* | -15, -28, 4 | 13, -27, 8 |
| | | Tha_L(R)_8_7 | 243 | 244 | *cTtha, caudal temporal thalamus* | -12, -22, 13 | 10, -14, 14 |
| | | Tha_L(R)_8_8 | 245 | 246 | *lPFtha, lateral pre-frontal thalamus* | -11, -14, 2 | 13, -16, 7 |

*Brain regions extracted in the first age-related dense subnetwork (*Orange color* )
**Brain regions extracted in the second age-related dense subnetwork (*Skyblue color* )

### Table S2b. Brain regions in the extracted subnetworks from HCP-A

| Lobe | Gyrus | Left and Right Hemisphere | Label ID.L | Label ID.R | Anatomical and modified Cyto-architectonic descriptions | lh.MNI(X,Y,Z) | rh.MNI(X,Y,Z) |
|---|---|---|---|---|---|---|---|
| Frontal Lobe | SFG, Superior Frontal Gyrus | SFG_L(R)_7_1 | 1 | 2 | *A8m, medial area 8* | -5 ,15, 54 | 7, 16, 54 |
| | | SFG_L(R)_7_2 | 3 | 4 | *A8dl, dorsolateral area 8* | -18, 24, 53 | 22, 26, 51 |
| | | SFG_L(R)_7_3 | 5 | 6 | *A9l, lateral area 9* | -11, 49, 40 | 13, 48, 40 |
| | | SFG_L(R)_7_4 | 7 | 8 | *A6dl, dorsolateral area 6* | -18, -1, 65 | 20, 4, 64 |
| | | SFG_L(R)_7_5 | 9 | 10 | *A6m, medial area 6* | -6, -5, 58 | 7, -4, 60 |
| | | SFG_L(R)_7_6 | 11 | 12 | *A9m,medial area 9* | -5, 36, 38 | 6, 38, 35 |
| | | SFG_L(R)_7_7 | 13 | 14 | *A10m, medial area 10* | -8, 56, 15 | 8, 58, 13 |
| | MFG, Middle Frontal Gyrus | MFG_L(R)_7_1 | 15 | 16 | *A9/46d, dorsal area 9/46* | -27, 43, 31 | 30, 37, 36 |
| | | MFG_L(R)_7_2 | 17 | 18 | *IFJ, inferior frontal junction* | -42, 13, 36 | 42, 11, 39 |
| | | MFG_L(R)_7_3 | 19 | 20 | *A46, area 46* | -28, 56, 12 | 28, 55, 17 |
| | | MFG_L(R)_7_4 | 21 | 22 | *A9/46v, ventral area 9/46* | -41, 41, 16 | 42, 44, 14 |
| | | MFG_L(R)_7_5 | 23 | 24 | *A8vl, ventrolateral area 8* | -33, 23, 45 | 42, 27, 39 |
| | | MFG_L(R)_7_6 | 25 | 26 | *A6vl, ventrolateral area 6* | -32, 4, 55 | 34, 8, 54 |
| | | MFG_L(R)_7_7 | 27 | 28 | *A10l, lateral area10* | -26, 60, -6 | 25, 61, -4 |
| | IFG, Inferior Frontal Gyrus | IFG_L(R)_6_1 | 29 | 30 | *A44d,dorsal area 44* | -46, 13, 24 | 45, 16, 25 |
| | | IFG_L(R)_6_2 | 31 | 32 | *IFS, inferior frontal sulcus* | -47, 32, 14 | 48, 35, 13 |
| | | IFG_L(R)_6_3 | 33 | 34 | *A45c, caudal area 45* | -53, 23, 11 | 54, 24, 12 |
| | | IFG_L(R)_6_4 | 35 | 36 | *A45r, rostral area 45* | -49, 36, -3 | 51, 36, -1 |
| | | IFG_L(R)_6_5 | 37 | 38 | *A44op, opercular area 44* | -39, 23, 4 | 42, 22, 3 |
| | | IFG_L(R)_6_6 | 39 | 40 | *A44v, ventral area 44* | -52, 13, 6 | 54, 14, 11 |
| | OrG, Orbital Gyrus | OrG_L(R)_6_1 | 41 | 42 | *A14m, medial area 14* | -7, 54, -7 | 6, 47, -7 |
| | | OrG_L(R)_6_2 | 43 | 44 | *A12/47o, orbital area 12/47* | -36, 33, -16 | 40, 39, -14 |
| | | OrG_L(R)_6_3 | 45 | 46 | *A11l, lateral area 11* | -23, 38, -18 | 23, 36, -18 |
| | | OrG_L(R)_6_4 | 47 | 48 | *A11m, medial area 11* | -6, 52, -19 | 6, 57, -16 |
| | | OrG_L(R)_6_5 | 49 | 50 | *A13, area 13* | -10, 18, -19 | 9, 20, -19 |
| | | OrG_L(R)_6_6 | 51 | 52 | *A12/47l, lateral area 12/47* | -41, 32, -9 | 42, 31, -9 |
| | PrG, Precentral Gyrus | PrG_L(R)_6_1 | 53 | 54 | *A4hf, area 4(head and face region)* | -49, -8, 39 | 55, -2, 33 |
| | | PrG_L(R)_6_2 | 55 | 56 | *A6cdl, caudal dorsolateral area 6* | -32, -9, 58 | 33, -7, 57 |
| | | PrG_L(R)_6_3 | 57 | 58 | *A4ul, area 4(upper limb region)* | -26, -25, 63 | 34, -19, 59 |
| | | PrG_L(R)_6_4 | 59 | 60 | *A4t, area 4(trunk region)* | -13, -20, 73 | 15, -22, 71 |
| | | PrG_L(R)_6_5 | 61 | 62 | *A4tl, area 4(tongue and larynx region)* | -52, 0, 8 | 54, 4, 9 |
| | | PrG_L(R)_6_6 | 63 | 64 | *A6cvl, caudal ventrolateral area 6* | -49, 5, 30 | 51, 7, 30 |
| | PCL, Paracentral Lobule | PCL_L(R)_2_1 | 65 | 66 | *A1/2/3ll, area1/2/3 (lower limb region)* | -8, -38, 58 | 10, -34, 54 |
| | | PCL_L(R)_2_2 | 67 | 68 | *A4ll, area 4, (lower limb region)* | -4, -23, 61 | 5, -21, 61 |
| Temporal Lobe | STG, Superior Temporal Gyrus | STG_L(R)_6_1 | 69 | 70 | *A38m, medial area 38* | -32, 14, -34 | 31, 15, -34 |
| | | STG_L(R)_6_2 | 71 | 72 | *A41/42, area 41/42* | -54, -32, 12 | 54, -24, 11 |
| | | STG_L(R)_6_3 | 73 | 74 | *TE1.0 and TE1.2* | -50, -11, 1 | 51, -4, -1 |
| | | STG_L(R)_6_4 | 75 | 76 | *A22c, caudal area 22* | -62, -33, 7 | 66, -20, 6 |
| | | STG_L(R)_6_5 | 77 | 78 | *A38l, lateral area 38* | -45, 11, -20 | 47, 12, -20 |
| | | STG_L(R)_6_6 | 79 | 80 | *A22r, rostral area 22* | -55, -3, -10 | 56, -12, -5 |
| | MTG, Middle Temporal Gyrus | MTG_L(R)_4_1 | 81 | 82 | *A21c, caudal area 21* | -65, -30, -12 | 65, -29, -13 |
| | | MTG_L(R)_4_2 | 83 | 84 | *A21r, rostral area 21* | -53, 2, -30 | 51, 6, -32 |
| | | MTG_L(R)_4_3 | 85 | 86 | *A37dl, dorsolateral area37* | -59, -58, 4 | 60, -53, 3 |
| | | MTG_L(R)_4_4 | 87 | 88 | *aSTS, anterior superior temporal sulcus* | -58, -20, -9 | 58, -16, -10 |
| | ITG, Inferior Temporal Gyrus | ITG_L(R)_7_1 | 89 | 90 | *A20iv, intermediate ventral area 20* | -45, -26, -27 | 46, -14, -33 |
| | | ITG_L(R)_7_2 | 91 | 92 | *A37elv, extreme lateroventral area37* | -51, -57, -15 | 53, -52, -18 |
| | | ITG_L(R)_7_3 | 93 | 94 | *A20r, rostral area 20* | -43, -2, -41 | 40, 0, -43 |
| | | ITG_L(R)_7_4 | 95 | 96 | *A20il, intermediate lateral area 20* | -56, -16, -28 | 55, -11, -32 |
| | | ITG_L(R)_7_5 | 97 | 98 | *A37vl, ventrolateral area 37* | -55, -60, -6 | 54, -57, -8 |
| | | ITG_L(R)_7_6 | 99 | 100 | *A20cl, caudolateral of area 20* | -59, -42, -16 | 61, -40, -17 |
| | | ITG_L(R)_7_7 | 101 | 102 | *A20cv, caudoventral of area 20* | -55, -31, -27 | 54, -31, -26 |
| | FuG, Fusiform Gyrus | FuG_L(R)_3_1 | 103 | 104 | *A20rv, rostroventral area 20* | -33, -16, -32 | 33, -15, -34 |
| | | FuG_L(R)_3_2 | 105 | 106 | *A37mv, medioventral area37* | -31, -64, -14 | 31, -62, -14 |
| | | FuG_L(R)_3_3 | 107 | 108 | *A37lv, lateroventral area37* | -42, -51, -17 | 43, -49, -19 |
| | PhG, Parahippocampal Gyrus | PhG_L(R)_6_1 | 109 | 110 | *A35/36r, rostral area 35/36* | -27, -7, -34 | 28, -8, -33 |
| | | PhG_L(R)_6_2 | 111 | 112 | *A35/36c, caudal area 35/36* | -25, -25, -26 | 26, -23, -27 |
| | | PhG_L(R)_6_3 | 113 | 114 | *TL, area TL (lateral PPHC, posterior parahippocampal gyrus)* | -28, -32, -18 | 30, -30, -18 |
| | | PhG_L(R)_6_4 | 115 | 116 | *A28/34, area 28/34 (EC, entorhinal cortex)* | -19, -12, -30 | 19, -10, -30 |
| | | PhG_L(R)_6_5 | 117 | 118 | *TI, area TI(temporal agranular insular cortex)* | -23, 2, -32 | 22, 1, -36 |
| | | PhG_L(R)_6_6 | 119 | 120 | *TH, area TH (medial PPHC)* | -17, -39, -10 | 19, -36, -11 |
| | pSTS, posterior Superior Temporal Sulcus | pSTS_L(R)_2_1 | 121 | 122 | *rpSTS, rostroposterior superior temporal sulcus* | -54, -40, 4 | 53, -37, 3 |
| | | pSTS_L(R)_2_2 | 123 | 124 | *cpSTS, caudoposterior superior temporal sulcus* | -52, -50, 11 | 57, -40, 12 |
| Parietal Lobe | SPL, Superior Parietal Lobule | SPL_L(R)_5_1 | 125 | 126 | *A7r, rostral area 7* | -16, -60, 63 | 19, -57, 65 |
| | | SPL_L(R)_5_2 | 127 | 128 | *A7c, caudal area 7* | -15, -71, 52 | 19, -69, 54 |
| | | SPL_L(R)_5_3 | 129 | 130 | *A5l, lateral area 5* | -33, -47, 50 | 35, -42, 54 |
| | | SPL_L(R)_5_4 | 131 | 132 | *A7pc, postcentral area 7* | -22, -47, 65 | 23, -43, 67 |
| | | SPL_L(R)_5_5 | 133 | 134 | *A7ip, intraparietal area 7(hIP3)* | -27, -59, 54 | 31, -54, 53 |
| | IPL, Inferior Parietal Lobule | IPL_L(R)_6_1 | 135 | 136 | *A39c, caudal area 39(PGp)* | -34, -80, 29 | 45, -71, 20 |
| | | IPL_L(R)_6_2 | 137 | 138 | *A39rd, rostrodorsal area 39(Hip3)* | -38, -61, 46 | 39, -65, 44 |
| | | IPL_L(R)_6_3 | 139 | 140 | *A40rd, rostrodorsal area 40(PFt)* | -51, -33, 42 | 47, -35, 45 |
| | | IPL_L(R)_6_4 | 141 | 142 | *A40c, caudal area 40(PFm)* | -56, -49, 38 | 57, -44, 38 |
| | | IPL_L(R)_6_5 | 143 | 144 | *A39rv, rostroventral area 39(PGa)* | -47, -65, 26 | 53, -54, 25 |
| | | IPL_L(R)_6_6 | 145 | 146 | *A40rv, rostroventral area 40(PFop)* | -53, -31, 23 | 55, -26, 26 |
| | Pcun, Precuneus | PCun_L(R)_4_1 | 147 | 148 | *A7m, medial area 7(PEp)* | -5, -63, 51 | 6, -65, 51 |
| | | PCun_L(R)_4_2 | 149 | 150 | *A5m, medial area 5(PEm)* | -8, -47, 57 | 7, -47, 58 |
| | | PCun_L(R)_4_3 | 151 | 152 | *dmPOS, dorsomedial parietooccipital sulcus(PEr)* | -12, -67, 25 | 16, -64, 25 |
| | | PCun_L(R)_4_4 | 153 | 154 | *A31, area 31 (Lc1)* | -6, -55, 34 | 6, -54, 35 |
| | PoG, Postcentral Gyrus | PoG_L(R)_4_1 | 155 | 156 | *A1/2/3ulhf, area 1/2/3(upper limb, head and face region)* | -50, -16, 43 | 50, -14, 44 |
| | | PoG_L(R)_4_2 | 157 | 158 | *A1/2/3tonIa, area 1/2/3(tongue and larynx region)* | -56, -14, 16 | 56, -10, 15 |
| | | PoG_L(R)_4_3 | 159 | 160 | *A2, area 2* | -46, -30, 50 | 48, -24, 48 |
| | | PoG_L(R)_4_4 | 161 | 162 | *A1/2/3tru, area1/2/3(trunk region)* | -21, -35, 68 | 20, -33, 69 |
| Insular Lobe | INS, Insular Gyrus | INS_L(R)_6_1 | 163 | 164 | *G, hypergranular insula* | -36, -20, 10 | 37, -18, 8 |
| | | INS_L(R)_6_2 | 165 | 166 | *vIa, ventral agranular insula* | -32, 14, -13 | 33, 14, -13 |
| | | INS_L(R)_6_3 | 167 | 168 | *dIa, dorsal agranular insula* | -34, 18, 1 | 36, 18, 1 |

| | | | | | | | |
|---|---|---|---|---|---|---|---|
| **Insular Lobe** | **INS, Insular Gyrus** | INS_L(R)_6_4 | 169 | 170 | *vId/vIg, ventral dysgranular and granular insula* | -38, -4, -9 | 39, -2, -9 |
| | | INS_L(R)_6_5 | 171 | 172 | *dIg, dorsal granular insula* | -38, -8, 8 | 39, -7, 8 |
| | | INS_L(R)_6_6 | 173 | 174 | *dId, dorsal dysgranular insula* | -38, 5, 5 | 38, 5, 5 |
| **Limbic Lobe** | **CG, Cingulate Gyrus** | CG_L(R)_7_1 | 175 | 176 | *A23d, dorsal area 23* | -4, -39, 31 | 4, -37, 32 |
| | | CG_L(R)_7_2 | 177 | 178 | *A24rv, rostroventral area 24* | -3, 8, 25 | 5, 22, 12 |
| | | CG_L(R)_7_3 | 179 | 180 | *A32p, pregenual area 32* | -6, 34, 21 | 5, 28, 27 |
| | | CG_L(R)_7_4 | 181 | 182 | *A23v, ventral area 23* | -8, -47, 10 | 9, -44, 11 |
| | | CG_L(R)_7_5 | 183 | 184 | *A24cd, caudodorsal area 24* | -5, 7, 37 | 4, 6, 38 |
| | | CG_L(R)_7_6 | 185 | 186 | *A23c, caudal area 23* | -7, -23, 41 | 6, -20, 40 |
| | | CG_L(R)_7_7 | 187 | 188 | *A32sg, subgenual area 32* | -4, 39, -2 | 5, 41, 6 |
| **Occipital Lobe** | **MVOcC, MedioVentral Occipital Cortex** | MVOcC _L(R)_5_1 | 189 | 190 | *cLinG, caudal lingual gyrus* | -11, -82, -11 | 10, -85, -9 |
| | | MVOcC _L(R)_5_2 | 191 | 192 | *rCunG, rostral cuneus gyrus* | -5, -81, 10 | 7, -76, 11 |
| | | MVOcC _L(R)_5_3 | 193 | 194 | *cCunG, caudal cuneus gyrus* | -6, -94, 1 | 8, -90, 12 |
| | | MVOcC _L(R)_5_4 | 195 | 196 | *rLinG, rostral lingual gyrus* | -17, -60, -6 | 18, -60, -7 |
| | | MVOcC _L(R)_5_5 | 197 | 198 | *vmPOS,ventromedial parietooccipital sulcus* | -13, -68, 12 | 15, -63, 12 |
| | **LOcC, lateral Occipital Cortex** | LOcC_L(R)_4_1 | 199 | 200 | *mOccG, middle occipital gyrus* | -31, -89, 11 | 34, -86, 11 |
| | | LOcC _L(R)_4_2 | 201 | 202 | *V5/MT+, area V5/MT+* | -46, -74, 3 | 48, -70, -1 |
| | | LOcC _L(R)_4_3 | 203 | 204 | *OPC, occipital polar cortex* | -18, -99, 2 | 22, -97, 4 |
| | | LOcC_L(R)_4_4 | 205 | 206 | *iOccG, inferior occipital gyrus* | -30, -88, -12 | 32, -85, -12 |
| | | LOcC _L(R)_2_1 | 207 | 208 | *msOccG, medial superior occipital gyrus* | -11, -88, 31 | 16, -85, 34 |
| | | LOcC _L(R)_2_2 | 209 | 210 | *lsOccG, lateral superior occipital gyrus* | -22, -77, 36 | 29, -75, 36 |
| **Subcortical Nuclei** | **Amyg, Amygdala** | Amyg_L(R)_2_1 | 211 | 212 | *mAmyg, medial amygdala* | -19, -2, -20 | 19, -2, -19 |
| | | Amyg_L(R)_2_2 | 213 | 214 | *lAmyg, lateral amygdala* | -27, -4, -20 | 28, -3, -20 |
| | **Hipp, Hippocampus** | Hipp_L(R)_2_1 | 215 | 216 | *rHipp, rostral hippocampus* | -22, -14, -19 | 22, -12, -20 |
| | | Hipp_L(R)_2_2 | 217 | 218 | *cHipp, caudal hippocampus* | -28, -30, -10 | 29, -27, -10 |
| | **BG, Basal Ganglia** | BG_L(R)_6_1 | 219 | 220 | *vCa, ventral caudate* | -12, 14, 0 | 15, 14, -2 |
| | | BG_L(R)_6_2 | 221 | 222 | *GP, globus pallidus* | -22, -2, 4 | 22, -2, 3 |
| | | BG_L(R)_6_3 | 223 | 224 | *NAC, nucleus accumbens* | -17, 3, -9 | 15, 8, -9 |
| | | BG_L(R)_6_4 | 225 | 226 | *vmPu, ventromedial putamen* | -23, 7, -4 | 22, 8, -1 |
| | | BG_L(R)_6_5 | 227 | 228 | *dCa, dorsal caudate* | -14, 2, 16 | 14, 5, 14 |
| | | BG_L(R)_6_6 | 229 | 230 | *dlPu, dorsolateral putamen* | -28, -5, 2 | 29, -3, 1 |
| | **Tha, Thalamus** | Tha_L(R)_8_1 | 231 | 232 | *mPFtha, medial pre-frontal thalamus* | -7, -12, 5 | 7, -11, 6 |
| | | Tha_L(R)_8_2 | 233 | 234 | *mPMtha, pre-motor thalamus* | -18, -13, 3 | 12, -14, 1 |
| | | Tha_L(R)_8_3 | 235 | 236 | *Stha, sensory thalamus* | -18, -23, 4 | 18, -22, 3 |
| | | Tha_L(R)_8_4 | 237 | 238 | *rTtha, rostral temporal thalamus* | -7, -14, 7 | 3, -13, 5 |
| | | Tha_L(R)_8_5 | 239 | 240 | *PPtha, posterior parietal thalamus* | -16, -24, 6 | 15, -25, 6 |
| | | Tha_L(R)_8_6 | 241 | 242 | *Otha, occipital thalamus* | -15, -28, 4 | 13, -27, 8 |
| | | Tha_L(R)_8_7 | 243 | 244 | *cTtha, caudal temporal thalamus* | -12, -22, 13 | 10, -14, 14 |
| | | Tha_L(R)_8_8 | 245 | 246 | *lPFtha, lateral pre-frontal thalamus* | -11, -14, 2 | 13, -16, 7 |

*Brain regions extracted in the first age-related dense subnetwork (*Orange color*)
**Brain regions extracted in the second age-related dense subnetwork (*Skyblue color*)

**Table S3. Quality control summary of cognitive data**

| UKB data field | Description | Cognitive test | Cognitive domain | Quality control | Transformation |
|---|---|---|---|---|---|
| 20023 | Mean time to correctly identify matches | Reaction time | Processing speed | Exclude if 1) accuracy rate < 50%; 2) hit rate <= 25%; 3) true negative rate <= 25%; or 4) total number of times snap-button pressed > 50 | Natural log (x) |
| 6373 | Number of puzzles correctly solved | Matrix pattern completion | Perceptual reasoning | Exclude if values < mean – 3SDs | NA |
| 399 | Number of incorrect matches in round | Paired matching | Visuospatial learning/memory | Exclude if number of correct matches in round < 6 | Natural log (x+1) |
| 6348 | Duration to complete numeric path (trail #1) | Trail making | Processing speed | Exclude if 1) did not complete trail or 2) made errors on trial #1 or trial #2 | 1) Calculate the difference between Trail #2 and Trail #1 by Trail #2 minus Trail #1; 2) Natural log(x) |
| 6350 | Duration to complete alphanumeric path (trail #2 | Trail making | Cognitive flexibility | | |
| 23324 | Number of symbol digit matches made correctly | Symbol digit substitution | Processing speed | Exclude if 1) less than 50% accuracy of attempted substitutions or 2) attempted less than 5 substitutions | NA |
| 21004 | Number of puzzles correct | Tower rearranging | Executive function/planning | Exclude if number of puzzles correct = 0 (may reflect failure to understand the task) | NA |
| 4282 | Maximum digits remembered correctly | Numeric memory | Working memory | Exclude if chose to abandon the test before completing the first round | NA |
| 20016 | Fluid intelligence score | Fluid intelligence/reasoning | Fluid intelligence | NA | NA |
